# Availability of an inflammatory macrophage niche drives phenotypic and functional alterations in Kupffer cells

**DOI:** 10.1101/2024.04.23.590829

**Authors:** Han-Ying Huang, Yan-Zhou Chen, Xin-Nan Zheng, Jia-Xing Yue, Huai-Qiang Ju, Yan-Xia Shi, Lin Tian

## Abstract

Inflammatory signals lead to recruitment of circulating monocytes and induce their differentiation into disease-associated macrophages^1–3^. Therefore, whether blocking inflammatory monocytes can mitigate disease progression is being actively evaluated^4^. Here, we employed multiple lineage tracing models and confirmed that monocyte-derived macrophages (mo-macs) are the major population of liver metastasis-associated macrophages (LMAMs), while the population of Kupffer cells (KCs), liver-resident macrophages, is diminished in liver metastatic nodules. Paradoxically, genetic ablation of mo-macs resulted in only a marginal decrease in LMAMs. Using a proliferation recording system and a KC tracing model in a monocyte-deficient background, we found that LMAMs can be replenished either via increased local macrophage proliferation or by promoting KC infiltration. After occupying macrophage niches left vacant by monocyte depletion, KCs exhibit substantial phenotypic and functional alterations through epigenetic reprogramming. These data suggest that dual blockade of monocytes and macrophages could be used to effectively target immunosuppressive myelopoiesis and to reprogram the microenvironment towards an immunostimulatory state.

## Main

Liver metastasis, the secondary malignancy frequently occurs in patients with gastrointestinal cancer and breast carcinoma, is an aggressive form of cancer progression with disappointing prognosis outcome and limited treatment options^5^. Immunotherapies have elicited durable treatment responses in some cancers but are ineffective in liver metastasis, which is enriched with various immune-suppressive cells and paralyzed effector and cytotoxic T cells^6,7^.

Tumor-associated hepatic myeloid cells, which consist of inflammatory monocytes and macrophages, are abundant in liver metastasis and foster a systemic immune desert microenvironment by inducing apoptosis of cytotoxic T cells^8^. In addition to T cell anergy, hepatic myeloid cells modulate microenvironment of metastasis through promoting angiogenesis, inducing fibrosis, remodeling extracellular matrix^9^. Therapeutic approaches that precisely targeting tumor-promoting myeloid cells could offer promise, but is hampered by a lack of knowledge about origin and functional maintenance of liver-metastasis associated macrophages (LMAMs).

In heathy liver, the self-renewal resident macrophages, also called Kupffer cells (KCs), line sinusoidal vessels and are in prime position to clear unfit red blood cells and even non-apoptotic disseminated tumor cells (DTCs) from portal vein^3^. The inflammatory milieu breaks the tissue homeostasis by disrupting KC niches and triggering an influx of blood monocytes and their differentiated macrophages^1,2^. However, the definitive KC tracing evidence is lacking so that it is difficult to delineate the two possibilities: 1) the replacement of KCs by mo-macs and 2) the phenotypic plasticity of KCs.

Here we dissected contribution of different myeloid lineages to the pool of LMAMs by mapping the fates of mo-macs and KCs. We found two distinct mechanisms – local proliferation and KC infiltration – that replenish LMAMs when the supply of circulating monocytes is blocked. After KCs infiltrate into metastatic nodules, the inflammatory cues could partially erase the ontologically epigenetic memory of KCs and reprogram KCs resemble to mo-macs. Our findings illuminate the resilience of LMAMs uopn monocyte blockade and the previously underappreciated plasticity of tissue-resident macrophages, highlighting the importance of blocking monocytes and differentiated macrophages simultaneously to target tumor-promoting myelopoiesis and to switch the metastatic microenvironment from immune-suppressive to immune-stimulatory.

## Results

### Phenotypic and functional disparities between normal liver macrophages and liver metastasis-associated macrophages

To study the differences in macrophage populations between liver metastasis tissues and normal liver tissues, we engrafted MC38 and E0771 cells – murine colorectal cancer cells and breast cancer cells, respectively – into syngeneic immunocompetent mice via portal vein injection to establish experimental metastasis models (**Fig. 1a**). We then performed multicolour flow cytometry (FC) to identify the immune cells that were differentially enriched in liver metastases. Consistent with the proposed inflammatory environment in malignant tissues^10^, the total number of CD45^+^ leukocytes was greater in metastatic nodules than in nearby normal liver tissues (**Fig. 1b**). Remarkably, the population of CD11b^low^F4/80^high^ resident-like macrophages was diminished, whereas CD11b^high^F4/80^high^ bone marrow-derived macrophages became the dominant macrophage population in liver metastatic nodules (**Fig. 1c**).

**Figure 1.**
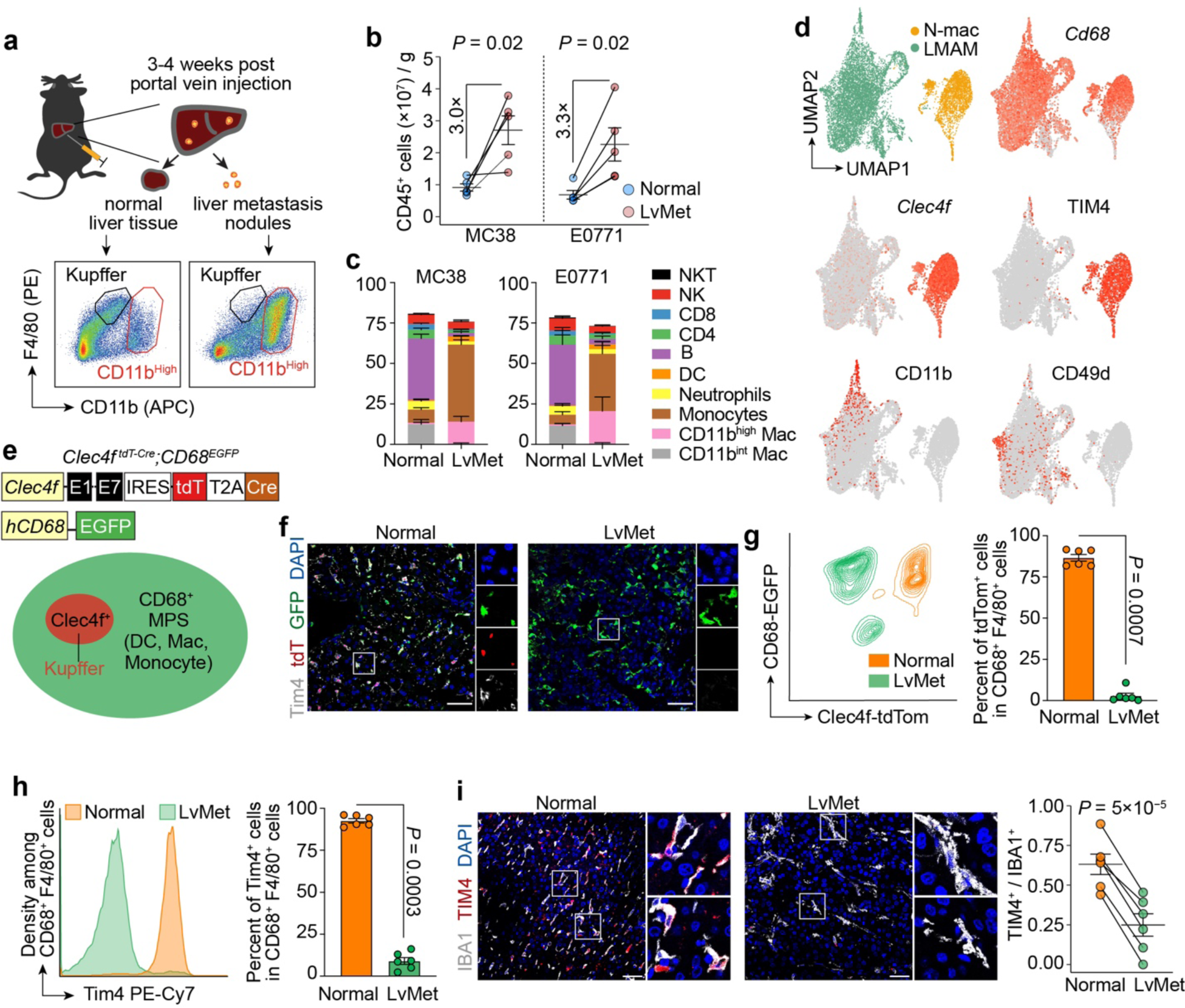
Phenotypic discrepancies between normal liver macrophages (N-macs) and liver metastasis-associated macrophages (LMAMs). **a-c).** Multi-colour flow cytometry analysis of immune cells in normal liver and liver metastasis (LvMet) (n=5 for MC38; n=5 for E0771). **d).** UMAP embedding of N-macs and LMAMs, and feature plots showing the expression of KC-identity genes and mo-mac signature genes. **e).** Schematic design of dual-fluorescent reporter mice for tracing macrophages. **f-h).** Representative immunofluorescence (IF) staining and flow cytometry quantification of KC markers Clec4f and Timd4 (n=6 for E0771; scale bar, 50 μm). **i).** Representative IF staining and quantification of Timd4 expression in liver metastasis tissue samples derived from breast cancer patients (triple negative, n=3; HER2^+^, n=3; scale bar, 50 μm). Mean ± s.e.m. shown. *P* values were calculated by comparing individual animals using two-tailed paired (**b**, **g**-**i**) Mann–Whitney *U*-test.

In addition to performing FC-based immunophenotyping, we also performed cellular indexing of transcriptomes and epitopes by sequencing (CITE-seq) on fluorescence-activated cell sorting (FACS)-sorted CD45^+^F4/80^+^ macrophages from metastatic nodules and nearby normal liver tissues, which allowed us to profile the transcriptome and 119 cell surface proteins simultaneously at the single-cell level. Both LMAMs and normal liver macrophages (N-macs) expressed *Cd68*, a marker of the mononuclear phagocyte system, which includes monocytes, macrophages, and dendritic cells. Monocyte-derived macrophage (mo-mac) markers such as Cd11b and Cd49d^11^ emerged on the cell surface of LMAMs (**Fig. 1d**). In contrast, the transcript level of *Clec4f*, a mouse KC marker, was substantially decreased in LMAMs (**Supplementary Videos 1, 2**), and this finding was confirmed by studies in dual-fluorescence *Clec4f ^tdT-Cre^;CD68^EGFP^* mice (**Fig. 1e**). In this model, the expression of the nuclear-localized red fluorescent protein tdTomato (tdT) was driven by the promoter of *Clec4f* ^12^, and the expression of enhanced green fluorescent protein (EGFP) was under the control of the promoter of *Cd68*^ref^^13^. Consistent with the CITE-seq results, N-macs were positive for both tdT and EGFP, while LMAMs were negative for tdT and expressed only EGFP (**Fig. 1f,g**).

Moreover, Timd4, the cell membrane protein that recognizes phosphoserine in the outer membrane leaflet of apoptotic cells, was not detected in LMAMs (**Fig. 1f,h**), and this finding was confirmed in liver metastases of breast cancer patients (**Fig. 1i**). When cocultured with apoptotic cancer cells labelled with CypHer5E, a dye that is weakly fluorescent at neutral pH but fluoresces brightly in phagosomes, tdT^+^EGFP^+^ N-macs but not tdT^−^EGFP^+^ LMAMs began to emit a strong far-red fluorescence signal within 1−2 hours (**Extended Data Fig. 1a-c**), suggesting that the loss of Timd4 prevented LMAMs from recognizing and engulfing apoptotic cells. To determine whether KCs can also engulf metastatic cancer cells *in vivo*, we also engrafted E0771 cancer cells engineered to express tdT into *wild-type* mice via portal vein injection (**Extended Data Fig. 1d**). Although Clec4f^+^ KCs could not effectively infiltrate metastatic tissues, the KCs engulfed nonapoptotic tdT^+^ cancer cells in the periphery of metastatic nodules (**Extended Data Fig. 1e**). Taken together, these results suggested that the macrophages in normal liver tissues and the macrophages in metastatic nodules displayed profoundly distinct morphologies, transcriptional profiles, and physiological functions.

### Liver metastasis-associated macrophages do not originate from Kupffer cells

Intrigued by the phenotypic discrepancies between N-macs and LMAMs, we next sought to identify the origin of LMAMs. Accumulating evidence suggests that monocytes are recruited to inflammatory sites^1^ and become the dominant population of LMAMs. This hypothetical model, however, does not rule out the possibility that KCs exhibit phenotypic plasticity through loss of their lineage markers and acquisition of mo-mac markers. To investigate the origin of LMAMs *in vivo*, we first crossed mice with KC-specific Cre recombinase expression (*Clec4f ^tdT-Cre^* mice) with the mice of the custom reporter line *R26^LSL-UPRT-HA-sfGFP^* (**Fig. 2a**). In the *Clec4f ^tdT-Cre^;R26^LSL-UPRT-HA-sfGFP^* mice, KCs and their progenies express superfolder green fluorescent protein (sfGFP) and HA-tagged uracil phosphoribosyl transferase (UPRT), which can be used to label nascent RNA specifically in KCs^14^. In particular, even when *Clec4f* is silenced (e.g., in LMAMs), cells derived from KCs retain the expression of the HA tag and sfGFP and can thus be distinguished from mo-macs, which never express *Clec4f*. According to both immunofluorescence (IF) staining and FC analysis, more than 90% of LMAMs did not express either the HA tag or sfGFP (**Fig. 2b,c**). To corroborate the findings in *Clec4f*-tracing mice, we also crossed sfGFP reporter mice with the mice expressing Cre recombinase in pre-macrophages from the yolk sac and embryo (*Tnfrsf11a^cre^*) (ref^15^) (**Extended Data Fig. 2a**). In this model, more than 80% of N-macs were positive for sfGFP, suggesting that fetal liver-derived KCs are maintained throughout the lifespan by self-renewal, with a minimal contribution from circulating monocytes. Similar to the findings in the KC tracing model using *Clec4f-Cre*, most LMAMs in this embryonic KC tracing model (*Tnfrsf11a^cre^;R26^LSL-UPRT-HA-sfGFP^*) were negative for the HA tag and sfGFP (**Extended Data Fig. 2b,c**), suggesting that KCs could not effectively infiltrate into metastatic tissues and contributed to the pool of LMAMs.

**Figure 2.**
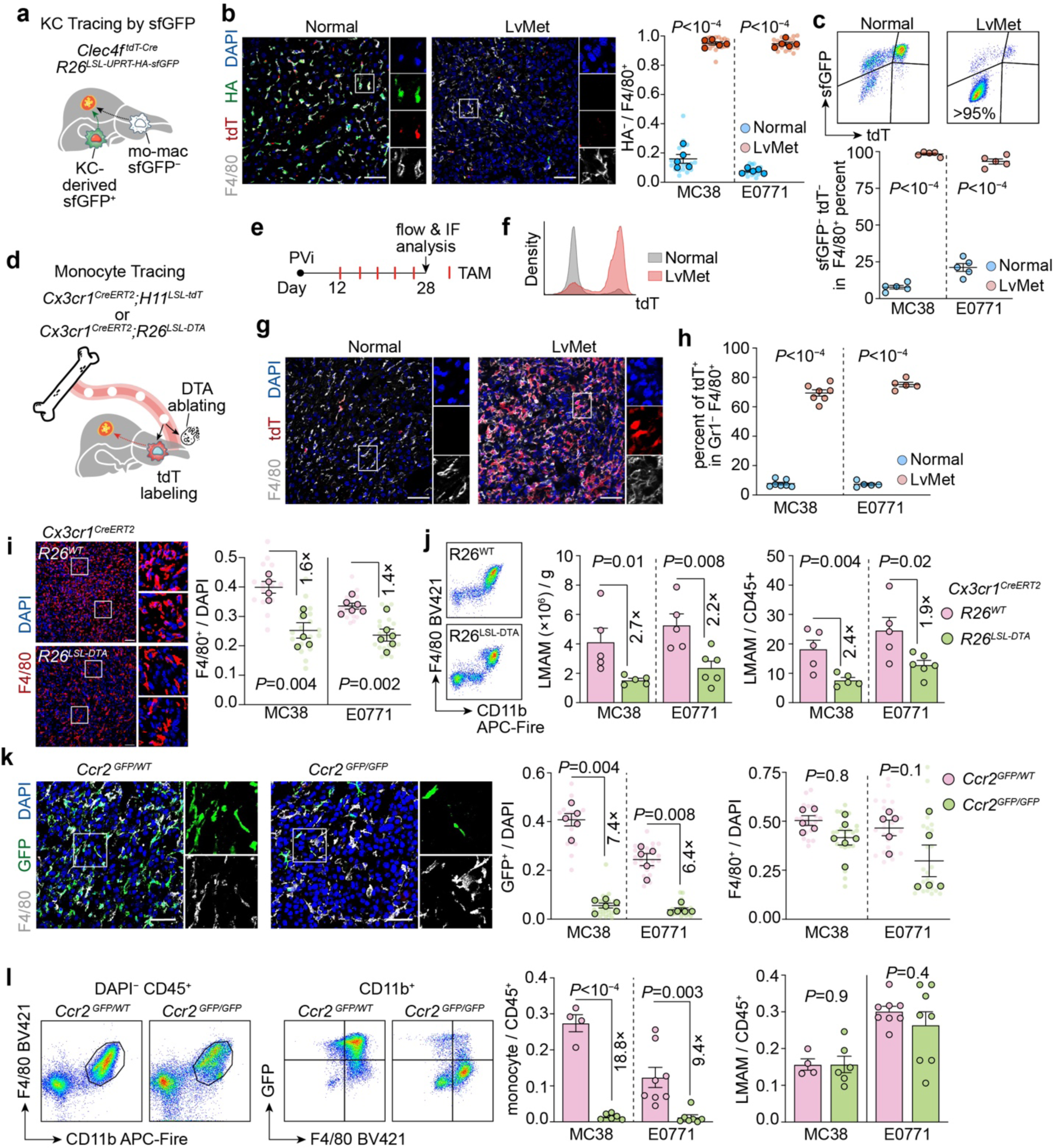
Fluorescent tracing and genetic ablation of distinct macrophage populations. **a).** Schematic design of KC lineage tracing mice. **b,c).** IF staining **(b)** and flow cytometry quantification **(c)** of KC-derived cells (HA tag^+^ or sfGFP^+^) (n=5 for MC38; n=5 for E0771; scale bar, 50 μm). **d,e).** Schematic and experimental design of monocyte lineage tracing and genetic ablation (IF, immunofluorescence). **f-h).** Representative histogram and IF quantification showing the expression of tdT in N-macs and LMAMs (n=7 for MC38; n=5 for E0771; scale bar, 50 μm). **i).** IF staining of LMAMs upon acute monocyte depletion (*Cx3cr1^CreERT^*^2^*;R26^LSL-DTA^*, n=5 for MC38, n=6 for E0771; littermate control *Cx3cr1^CreERT^*^2^*;R26^WT^*, n=4 for MC38; n=6 for E0771; scale bar, 50 μm). **j)** Flow cytometry quantification of LMAMs upon acute monocyte depletion (*Cx3cr1^CreERT^*^2^*;R26^LSL-DTA^*, n=5 for MC38, n=6 for E0771; littermate control *Cx3cr1^CreERT^*^2^*;R26^WT^*, n=5 for MC38; n=5 for E0771). **k).** IF quantification of monocytes and LMAMs by impairing monocyte chemotaxis (*Ccr2^GFP/GFP^*: n=6 for MC38; n=5 for E0771; *Ccr2^GFP/WT^* littermates: n=5 for MC38; n=5 for E0771; scale bar, 50 μm). **l).** Representative flow cytometry plot showing loss of Ccr2 expression in LMAMs of *Ccr2^GFP/GFP^*mice, and quantification of monocytes and LMAMs (*Ccr2^GFP/GFP^*: n=6 for MC38; n=8 for E0771; *Ccr2^GFP/WT^* littermates: n=4 for MC38; n=8 for E0771). Mean ± s.e.m. shown. Smaller dots are values from individual fields. Outlined circles are mean values taken over multiple fields/sections from the same mouse. *P* values were calculated by comparing individual animals using two-tailed paired (**b**, **c**, **h**) and unpaired (**i**-**l**) Mann–Whitney *U*-test.

Given that KCs reside in the liver and do not enter the circulation, we also performed a time-course parabiosis experiment (**Extended Data Fig. 2d**). After two weeks of blood circulation sharing between *CD45.2^+/+^;Tnfrsf11a^cre^;R26^LSL-UPRT-HA-sfGFP^*and *CD45.1^+/+^;CD68-EGFP* parabionts, we injected cancer cells into the portal vein of *CD45.1^+/+^*mice and waited another four weeks for the establishment of overt metastatic colonies in these mice. Indeed, very few LMAMs were positive for the HA tag (**Extended Data Fig. 2e**), and the majority of CD45.2^+^ LMAMs were negative for sfGFP (**Extended Data Fig. 2f**), supporting the idea that the LMAM population is composed mostly of macrophages derived from blood monocytes.

### Liver metastasis-associated macrophages are derived from monocytes, but can be replaced from other sources

In addition to tracing KCs, we also used an orthogonal approach of mapping the fate of inflammatory monocytes (*Cx3cr1^CreERT^*^2^*;H11^LSL-tdT^* for fluorescence tracing and *Cx3cr1^CreERT^*^2^*;R26^LSL-DTA^*for genetic ablation) (**Fig. 2d**). KCs are established prior to birth and are independent of Cx3cr1^ref^^16^; therefore, the fluorescent labelling model *Cx3cr1^CreERT^*^2^*;H11^LSL-tdT^*and genetic ablation model *Cx3cr1^CreERT^*^2^*; R26^LSL-DTA^* were useful for tracing and depleting monocytes, respectively, in a controllable manner via the oral administration of tamoxifen (**Fig. 2e**). Although more than 75% of LMAMs were positive for tdT, indicating that they were derived from monocytes (**Fig. 2f-h**), diphtheria toxin A (DTA)-mediated depletion of inflammatory monocytes only decreased the LMAM population by half (**Fig. 2i,j**). In a complementary approach, we employed Ccr2 knockout (*Ccr2^GFP/GFP^*) mice, in which the coding sequence of the chemokine receptor Ccr2 was replaced by that of GFP^17^. Notably, the protein level of Ccl2, the ligand of Ccr2, was markedly increased in the small liver metastatic lesions (**Extended Data Fig. 3**), indicating the potential role of the Ccl2-Ccr2 axis in monocyte recruitment. Similar to the aforementioned DTA-based ablation of monocytes, knockout of Ccr2 resulted in profound loss of monocytes (by 85% as determined by IF staining and by 90% as determined by FC) but only a marginal decrease in LMAMs (by 37% as determined by IF staining and 35% as determined by FC) (**Fig. 2k,l**). Collectively, these data indicate that the LMAM population is derived mostly from monocytes but may be partially replenished via other mechanisms when monocyte recruitment is blocked.

### Enhanced *in situ* macrophage proliferation upon monocyte blockade

To explain how LMAMs are replenished upon monocyte depletion, we proposed two possibilities: i) increased proliferation of local macrophages within metastatic nodules and ii) infiltration of KCs from nearby normal tissues. In fact, the proliferation of local macrophages or local monocytes to replenish vacant macrophage niches has been implied to occur in various malignant^18^ and normal tissues^19–21^ via colony-stimulating factor 1 (Csf1) and its receptor Csf1r. To determine whether local proliferation contributes to the replenishment of LMAMs, we established a proliferation recorder mouse model (*H11^DreERT^*^2^*;Ki67^RSR-Cre^;R26^LSL-UPRT-HA-sfGFP^* mice) (**Fig. 3a**), which was similar to the ProTracer system previously reported^22^. Unlike the method of identifying proliferating cells by IF staining for the cell proliferation marker Ki67, in this proliferation recorder system, cells that have undergone proliferation starting from a defined time point can be marked. Upon treatment with one tamoxifen pulse, DreERT2 excises the rox-flanked transcriptional termination signal upstream of the Cre recombinase sequence, yielding a new Ki67-Cre genotype in which Cre recombinase can label proliferating cells at any time thereafter. Using this proliferation recording tool, we found that only approximately 30% of LMAMs were sfGFP^+^, whereas more than 85% of tumor-associated endothelial cells were sfGFP^+^ (**Fig. 3b**). We then leveraged clodronate liposomes (CLs) to block the external supply of LMAMs. Although CLs can effectively deplete both circulating monocytes and KCs (**Fig. 3c**), they cannot cross the tumor vasculature due to their size; thus, macrophages within metastatic nodules are not depleted (**Fig. 3d,e**). Intravenous administration of CLs increased the percentage of sfGFP^+^ LMAMs by twofold compared to that in mice treated with control liposomes (**Fig. 3f-h**), suggesting that the local proliferation of LMAMs increased in response to monocyte depletion (**Fig. 3i**).

**Figure 3.**
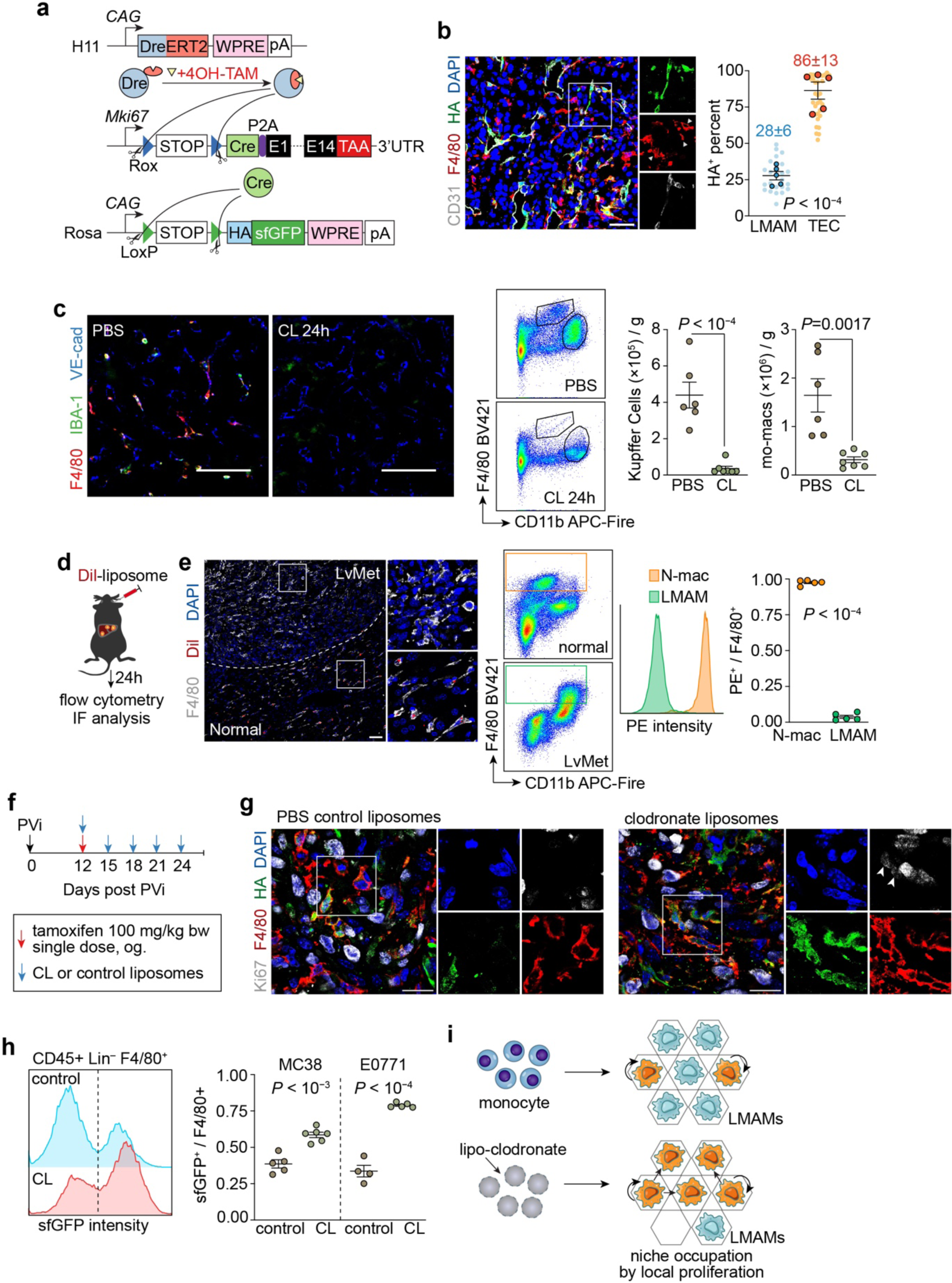
Enhanced local macrophage proliferation within liver metastasis upon monocyte depletion. **a).** Schematic design of the proliferation recorder mice (*H11^DreERT^*^2^*;Ki67-RSR-Cre; R26^LSL-UPRT-HA-sfGFP^*). **b).** Representative IF staining for HA tag, macrophage marker F4/80 and endothelial marker CD31 and quantification of fraction of proliferated cells in LMAMs and tumor-associated endothelial cells (TECs) (n=5; scale bar, 50 μm). **c).** Representative IF staining (scale bar, 50 μm) and flow cytometry quantification of Kupffer cells and mo-macs 24 h after intravenous injection of PBS control liposomes (n=6) or clodronate liposomes (CL, n=7). **d).** Schematic of labelling macrophages with red fluorescent dye Dil-loaded liposomes. **e).** Representative IF staining, and flow cytometry gating and quantification showing N-macs but not LMAMs engulfing Dil-loaded liposomes (n=5; scale bar, 50 μm). **f).** Schematic of experimental design. Metastasis-bearing proliferation recorder mice were pulsed with a single tamoxifen gavage, and were treated with CL or control liposomes every three days until the experimental endpoint (bw, body weight; og, oral gavage). **g).** Representative IF staining of HA tag^+^ proliferated LMAMs and Ki67^+^ proliferating LMAMs (indicated by white arrowheads; scale bar, 50 μm). **h).** Flow cytometry quantification of sfGFP^+^ proliferated LMAMs in CL-treated mice (n=6 for MC38, n=5 for E0771) and control mice (n=5 for MC38; n=4 for E0771). **i).** Schematic showing the activation of local macrophage proliferation upon blockade of monocyte supply. Mean ± s.e.m. shown. *P* values were calculated by comparing individual animals using two-tailed paired (**b**) and unpaired (**c**, **e**, **h**) Student’s *t*-test.

### Phenotypic plasticity of KCs

In addition to our hypothesis regarding the local proliferation of LMAMs, we hypothesized that KCs could also infiltrate liver metastases when the macrophage niche was available due to monocyte deficiency. To test this hypothesis, we combined the KC tracing system with Ccr2 knockout mice (*Clec4f ^tdT-Cre^*;*R26^LSL-UPRT-HA-sfGFP^;Ccr2^GFP/GFP^*). Because both Ccr2^+^ mo-macs and KC-derived LMAMs are positive for GFP, the GFP signal cannot be used to distinguish KC-derived cells from mo-macs. Thus, in addition to containing sfGFP and the HA tag, our Cre-inducible reporter line was also designed to incorporate a woodchuck hepatitis virus posttranscriptional regulatory element (WPRE) initially to increase mRNA transcript stability and expression; however, this element may also be used as a DNA tag to trace Cre recombinase-derived cells in single-cell RNA sequencing (scRNA-seq) analysis, because the WPRE is located at the 3’ end of the transcript upstream of the polyA signal (**Fig. 4a**). To test whether the WPRE can be detected in sfGFP^+^ cells, we independently crossed NK-specific Cre model mice^23^ with our reporter mice (*Ncr1^Cre^*; *R26^LSL-UPRT-HA-sfGFP^*) and sorted two NK1.1^+^ populations, namely, sfGFP^+^ NK cells and sfGFP^−^ NKT cells, by FACS for single-cell TCR sequencing (scTCR-seq) (**Extended Data Fig. 4a-c**). In both normal and metastatic tissues, greater numbers of WPRE^+^ cells were detected in the cluster of sfGFP^+^ TCR^−^ NK cells than in the cluster of sfGFP^−^ TCR^+^ NKT cells (**Extended Data Fig. 4d-f**), suggesting that the WPRE is useful as a heritable cell tag when fluorescent protein-based tracing is not feasible.

**Figure 4.**
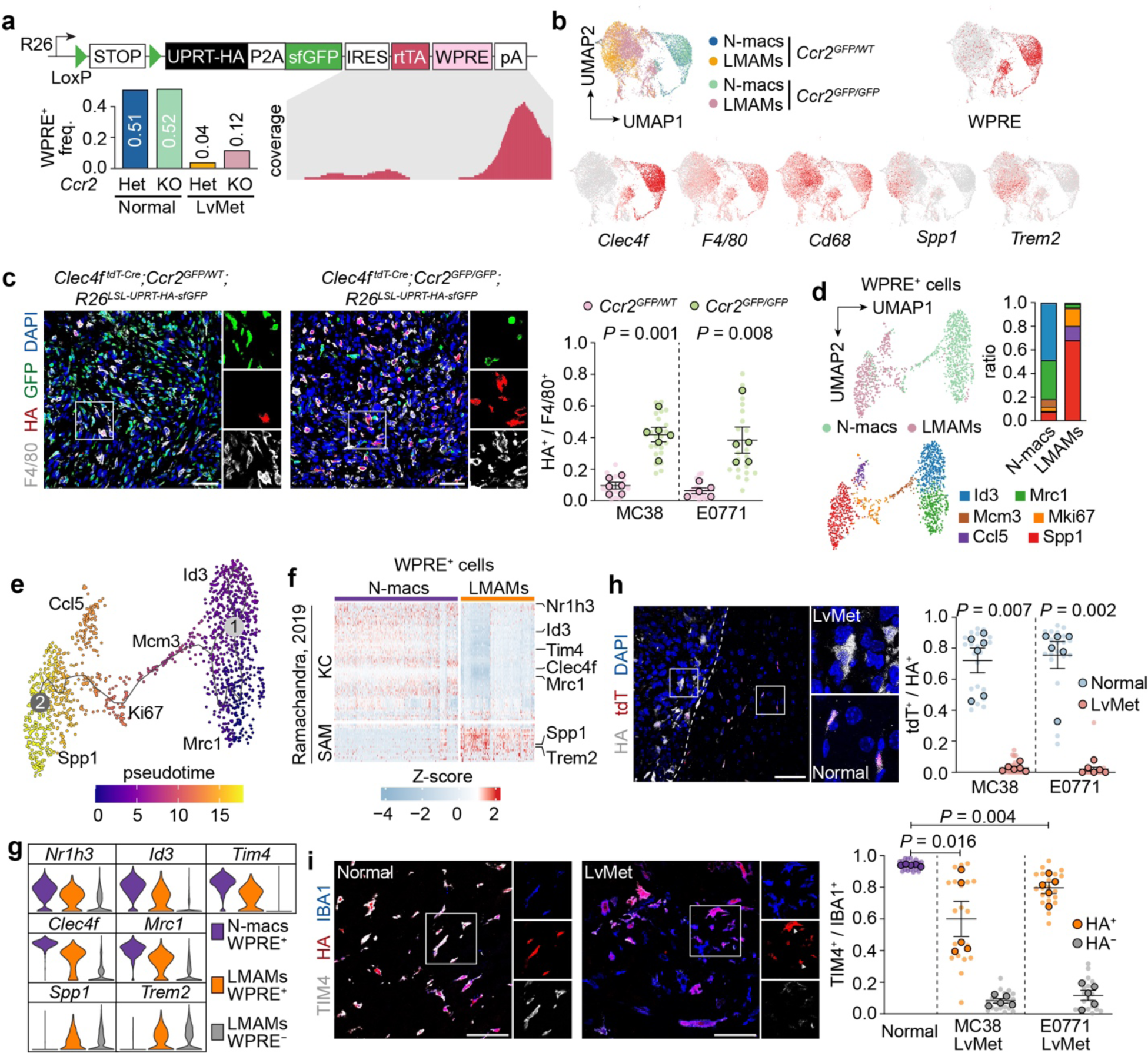
Phenotypic plasticity of KCs upon occupying mo-mac niche in liver metastasis. **a).** Bar plot showing the frequency of WPRE^+^ macrophages from different genetic background, and integrated genome viewer plot showing transcripts mapping to the WPRE. **b).** UMAP embedding of N-macs and LMAMs from *Ccr2^GFP/GFP^* mice and control littermates. Feature plots showing the expression of KC-identity genes and SAM genes. **c).** IF staining and quantification of HA tag^+^ KC-derived LMAMs (*Ccr2^GFP/GFP^*: n=6 for MC38; n=5 for E0771; *Ccr2^GFP/WT^* littermates: n=6 for MC38; n=5 for E0771). **d,e).** Re-clustering of WPRE^+^ N-macs and WPRE^+^ LMAMs from *Ccr2^GFP/GFP^* mice (**d**), and pseudo time analysis revealing the dynamic gene regulatory programs along of KC trans-differentiation (**e**). **f).** Heatmap showing the downregulation of KC-identity genes and upregulation of SAM genes in KCs when infiltrating into liver metastasis. **g).** Violin plots showing the expression of selected KC-identity genes and SAM genes in normal liver KCs (WPRE^+^ N-macs), KC-derived LMAMs (WPRE^+^ LMAMs) and mo-macs (WPRE^+^ LMAMs). **h,i**) IF staining and quantification of Clec4f expression (indicated by the nuclear-localized tdT) (**h**, n=6 for MC38; n=6 for E0771) and Timd4 expression (**i**, n=5 for MC38; n=5 for E0771). Mean ± s.e.m. shown. Smaller dots are values from individual fields. Outlined circles are mean values taken over multiple fields/sections from the same mouse. *P* values were calculated by comparing individual animals using two-tailed unpaired (**c,i**) and paired (**h**) Mann–Whitney *U*-test.

We then sorted N-macs and LMAMs isolated from *Clec4f ^tdT-Cre^*;*R26^LSL-UPRT-HA-sfGFP^;Ccr2^GFP/GFP^*mice and their *Ccr2^GFP/WT^* littermates by FACS and performed scRNA-seq to investigate the contribution of KCs to the pool of LMAMs. After aligning the sequencing reads to the transgenic reporter allele, we observed a significant peak over the WPRE and enrichment of WPRE^+^ cells in the N-mac population (**Fig. 4a,b**). A greater percentage of WPRE^+^ LMAMs was detected in the *Ccr2^GFP/GFP^* mice than in their *Ccr2^GFP/WT^* littermates (**Fig. 4a**). This finding was confirmed by IF staining for the HA tag (**Fig. 4c**), suggesting that KCs may occupy inflammatory macrophage niches left vacant by monocyte depletion. To understand the phenotypic plasticity of KCs during their infiltration into metastatic nodules, we performed pseudotime analysis on WPRE^+^ cells in our scRNA-seq dataset and found that Clec4f^+^ KCs in the normal liver (Id3-cluster and Mrc1-cluster) could transdifferentiate into Clec4f^−^ KCs (Spp1-cluster) via a status of increased proliferation (Mcm3 cluster and Mki67 cluster) (**Fig. 4d,e**). To further confirm that the KCs underwent cell division during transdifferentiation, we leveraged the HA-tagged UPRT protein in our reporter mice to profile the KC-specific transcriptome *in situ*^14^ (**Extended Data Fig.5a-e**) and performed IF staining for the HA tag and the cell proliferation marker Ki67 (**Extended Data Fig.5f**). Of particular interest, these KC-derived LMAMs, similar to mo-macs, expressed high levels of scar-associated macrophage (SAM) genes (e.g., Spp1 and Trem2) that were previously identified in macrophages from fibrotic livers^24^ (**Fig. 4f,g**) and exhibited downregulation of KC marker genes, including Clec4f and Timd4, and these changes were also associated with the morphological switch from an elongated shape to a flat shape (**Fig. 4h,i**).

### Functional alteration and epigenetic reprogramming of KCs

We therefore sought to determine whether this phenotypic plasticity of KCs can affect their functions. We generated *H11^DreERT^*^2^*;Clec4f ^tdT-Cre^;R26^LSL-RSR-tdT-DTR^*mice to conditionally label or ablate KCs (**Fig. 5a**). In this model, the expression of CAG promoter-driven cytoplasmic tdT and DTR can be activated by the oral administration of tamoxifen, allowing the tracing of KC-derived LMAMs when Clec4f-driven expression of nuclear tdT is repressed in liver metastasis (**Fig. 5b**). Consistent with the previously proposed anti-tumor effect of KCs^25–28^, depletion of KCs by intraperitoneal administration of diphtheria toxin (DT) increased the liver metastasis burden compared with that in mice treated with PBS (**Fig. 5b-d**). In addition to acute genetic ablation, both pharmacological depletion of KCs by CL treatment (**Extended Data Fig. 6a-d**) and genetic impairment of KCs by knockout of Id3 in embryonic KCs (**Extended Data Fig. 6e,f**) enhanced liver colonization, suggesting the potent tumor elimination effect of KCs.

**Figure 5.**
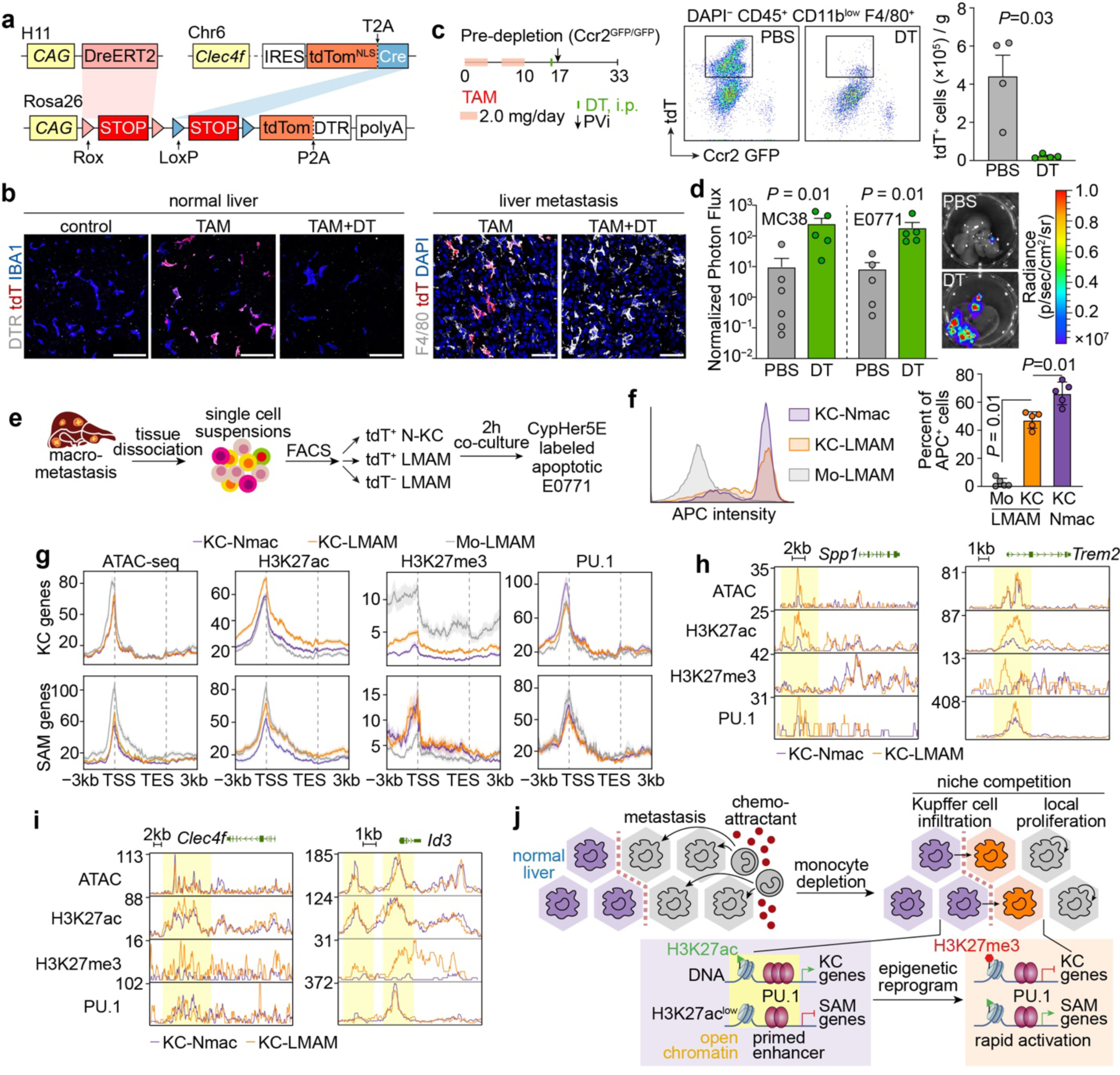
Functional alteration and epigenetic reprogramming of KCs in liver metastasis. **a).** Schematic design of KC-tdT-DTR mice (*H11^DreERT^*^2^*;Clec4f ^tdT-Cre^;R26^LSL-RSR-tdT-DTR^*) for tracing and ablating of KCs in normal tissues and metastatic tissues. **b).** Acute depletion of normal liver KCs (**b**) and KC-derived LMAMs (**c**) by activation of DreERT2 using tamoxifen (TAM) and then by intraperitoneally (i.p.) administration of diphtheria toxin (DT) (scale bar, 50 μm). **c).** Schematic diagram of pre-depletion of KCs and establishment of experimental liver metastasis models (PVi, portal vein injection). Flow cytometry quantification showing DT-mediated genetic ablation of tdT^+^ KCs (n=4 for PBS control group; n=4 for DT-treated group). **d).** Bar plots comparing liver metastasis burden of DT-treated mice (n=5 for MC38; n=5 for E0771) or PBS-treated control mice (n=6 for MC38; n=5 for E0771). The *ex vivo* bioluminescence value was normalised to the *in vivo* bioluminescence value obtained immediately after portal vein injection (day 0). **e).** Schematic diagram of enrichment of macrophages for in vitro co-culture with Staurosporine-treated apoptotic E0771 cells labelled with CypHer5E. **f).** Flow cytometry quantification of distinct macrophage populations phagocyting apoptotic cancer cells (n=5). **g).** Normalised distribution of ATAC-seq or CUT&Tag-seq densities from 3 kb upstream to transcription start sites (TSS) to 3 kb downstream to transcription end sites (TES) of 118 KC-identity genes or 46 SAM genes. **h,i**). Genome browser tracks of ATAC-seq or CUT&Tag-seq for H3K27ac, H3K27me3 and PU.1 signals in the vicinities of selected SAM genes (**h**) and KC-identity genes (**i**). The enhancer regions are highlighted in yellow. **j).** Mechanistic model showing the epigenetic reprogramming that underlies phenotypic and functional alterations of KCs in LvMet. Mean ± s.e.m. shown. *P* values were calculated by comparing individual animals using two-tailed unpaired (**d**) and paired (**f**) Mann–Whitney *U*-test.

We then sought to determine whether the loss of Timd4 in KC-derived LMAMs prevents them from engulfing apoptotic cells. When KCs enriched from normal and metastatic tissues by anti-F4/80 magnetic beads were cocultured with CypHer5E-labelled apoptotic cancer cells (**Fig. 5e**), KC-derived LMAMs exhibited impaired phagocytic activity compared with that of KCs in normal liver tissues but were still markedly different from mo-macs (**Fig. 5f**). Indeed, even though the expression level of Timd4 was markedly reduced, more than half of the KC-derived LMAMs were still positive for Timd4 (**Fig. 4i**). These data suggest that both niche factors and cell ontogeny factors may cooperatively control the phenotypes and functions of KCs.

To investigate the molecular mechanisms underlying the abovementioned phenotypic and functional alterations, we sorted tdT^+^ KCs from normal liver tissue and tdT^+^ KCs and tdT^−^ mo-macs from metastatic nodules by FACS and performed assay for transposase-accessible chromatin with sequencing (ATAC-seq) to identify open chromatin and cleavage under targets and tagmentation (CUT&Tag) to evaluate the occupancy of the macrophage-specific transcription factor PU.1 and histone modifications, including H3K27ac and H3K27me3^ref^^29^.

Surprisingly, the enhancers of most SAM genes (e.g., *Spp1* and *Trem2*) were accessible even in KCs in the normal liver and were poised for rapid activation via the acquisition of the transcriptional activation marker H3K27ac in response to metastatic niche factors (**Fig. 5g,h**). Furthermore, although KC-derived LMAMs inherited a cell ontogeny-imprinted enhancer landscape by maintaining open chromatin in KC marker genes, KC marker genes (e.g., *Clec4f* and *Id3*) could be silenced by the deposition of the transcriptional repression mark H3K27me3 (**Fig. 5i**). Together, these data suggested that the induction of SAM gene expression and the loss of KC marker gene expression in KC-derived LMAMs were attributed to epigenetic reprogramming in enhancer landscapes (**Fig. 5j**).

### Dual blockade of monocyte recruitment and macrophage proliferation reduces LMAMs more effectively

As local macrophage proliferation and KC infiltration compensated for monocyte deficiency, we proposed that dual blockade of monocyte recruitment and macrophage proliferation by knocking out Ccr2 and targeting Csf1r, respectively, can enhance the efficiency of LMAM depletion. Compared with knockout of Ccr2 alone or treatment with the Csf1r inhibitor PLX5622^ref30^ alone, dual inhibition resulted in more complete reduction of metastasis-associated monocytes and macrophages (**Fig. 6a-d**). Importantly, we found that this combination strategy could also be applied to subcutaneous MC38 tumors and orthotopic E0771 tumors, as the population of IBA1^+^ tumor-associated macrophages became negligible when Ccr2 and Csf1r were simultaneously blocked (**Extended Data Fig.7**).

**Figure 6.**
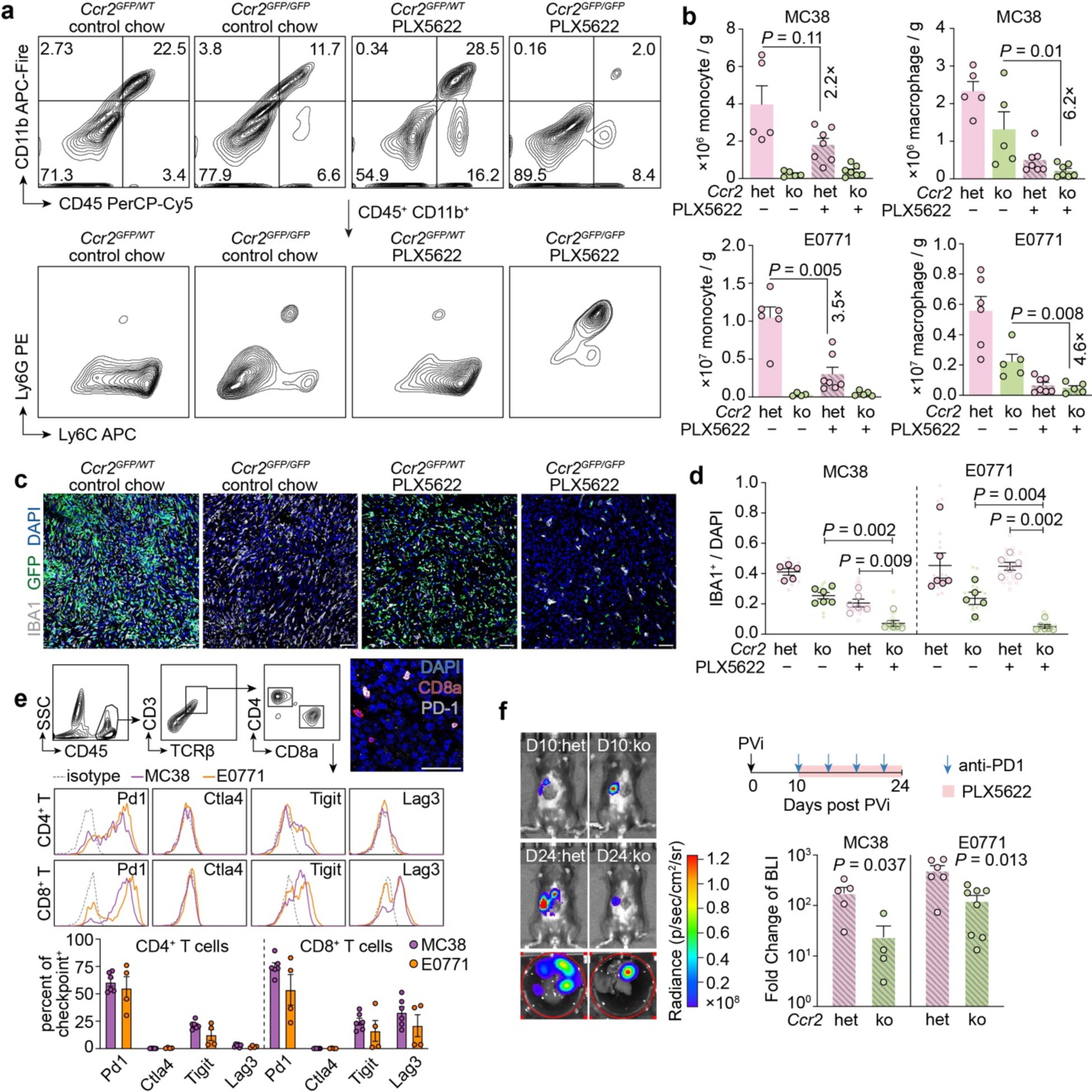
Increased efficacy of immune checkpoint inhibitor by dual blockade of monocyte infiltration and macrophage proliferation. **a,b).** Representative flow cytometry plots and quantification of LvMet infiltrating monocytes and macrophages in *Ccr2^GFP/GFP^* mice fed with PLX5622 diet (n=7 for MC38; n=5 for E0771) or control chow (n=5 for MC38; n=5 for E0771), and in *Ccr2^GFP/WT^*littermates fed with Csf1r inhibitor PLX5622 diet (n=7 for MC38; n=7 for E0771) or control chow (n=5 for MC38; n=6 for E0771). **c,d).** Representative IF staining and quantification for LvMet infiltrating IBA1^+^ myeloid cells in *Ccr2^GFP/GFP^* mice fed with PLX5622 diet (n=6 for MC38; n=6 for E0771) or control chow (n=6 for MC38; n=5 for E0771; scale bar, 50 μm), and in *Ccr2^GFP/WT^* littermates fed with Csf1r inhibitor PLX5622 diet (n=6 for MC38; n=6 for E0771) or control chow (n=5 for MC38; n=6 for E0771). **e).** Flow cytometry quantification of immune checkpoints in CD4^+^ T cells and CD8^+^ T cells from LvMet (n=6 for MC38; n=4 for E0771). Representative IF staining showing CD8^+^ T cells expressing Pd1 (scale bar, 50 μm). **f).** Representative bioluminescence imaging of metastasis-bearing *Ccr2^GFP/GFP^* mice and *Ccr2^GFP/WT^* littermates treated with PLX5622 and anti-Pd1 antibody (left). Bar plots comparing the effects of dual blockade of Pd1 and Csf1r on metastatic outgrowth of *Ccr2^GFP/GFP^* mice (n=4 for MC38; n=8 for E0771) or *Ccr2^GFP/WT^* littermates (n=5 for MC38; n=6 for E0771). The *ex vivo* bioluminescence value was normalised to the *in vivo* bioluminescence value of micrometastasis (day 10 post portal vein injection). Mean ± s.e.m. shown. *P* values were calculated by comparing individual animals using two-tailed unpaired Mann–Whitney *U*-test (**b**, **d**, **f**).

One important immunosuppressive role of LMAMs is to induce apoptosis in cytotoxic T cells^8^. Indeed, more than half of the CD4^+^ T cells and CD8^+^ T cells infiltrating liver metastatic nodules were positive for the immune checkpoint molecule programmed death-1 (Pd1) (**Fig. 6e**). In agreement with the findings of a previous study in which radiotherapy was used to eliminate LMAMs^8^, dual blockade of Ccr2 and Csf1r increased the efficacy of anti-Pd1 immune checkpoint inhibition compared with that achieved by knockout of Ccr2 alone (**Fig. 6f**).

## Discussion

The present work reveals the heterogeneity and plasticity of macrophages in liver metastasis. Compared with other liver diseases such as non-alcoholic fatty liver or steatohepatitis^31,32^, liver metastasis is almost devoid of KCs, and is more enriched with mo-macs, which is consistent with the niche competition model in inflammation^33^. Counterintuitively, depletion of monocytes does not lead to a significant reduction in LMAM numbers, either due to an enhanced proliferation of macrophages within the malignancy or due to the infiltration of KCs from nearby normal liver. Our results provide direct evidence for the niche competition model on the basis of monocyte deficiency, and revealed that at least a subpopulation of KCs can adapt to the inflammatory microenvironment through transient proliferation and subsequently trans-differentiate toward a cellular state similar to mo-macs.

Notably, the plasticity of tissue-resident macrophages is underappreciated, as the presence of mo-macs contaminates the resident macrophage pool^34^. In this study, we leveraged precise fate mapping of resident macrophage in the monocyte deficient background to uncover the plasticity of KCs in liver metastasis, which warrants further attention in other physiological and pathological contexts.

Of particular interest, both KC-derived and monocyte-derived LMAMs resemble SAMs previously identified in human and mouse liver cirrhosis^24^. The emergence of SAM genes and the loss of KC identity genes are attribute to the epigenetic landscape reprogramming. For example, the enhancers of SAM genes are always accessible even in KCs of normal liver and are poised for rapid activation by acquiring histone modification H3K27ac in response to metastasis niche factors. Furthermore, although KC-derived LMAMs inherit a cell ontogeny-imprinted enhancer landscape by maintaining open chromatin for KC-identity genes, the KC-identity genes can be silenced by the recruitment of histone modification H3K27me3.

Overall, our study points that cell ontogeny and enviromental factors jointly shape the epigenetic landscape, which further alters phenotpic plasticity and specifies physiological function. Because the replenishment from fully differentiated macrophages compensate for monocyte blockade, combinatory interventions perturbing both monocytes and macrophages represent a more effective approach to targeting tumor-promoting myeloid cells, and present promosing stratesies for other potential synergy.

## Methods

### Cell lines and cell culture

Mouse triple negative breast cancer (TNBC) E0771 cells (CH3 Biosystems) and mouse colorectal cancer MC38 cells (gift from Joan Massagué) were cultured in RPMI-1640 medium (ThermoFisher) and DMEM/High Glucose medium (ThermoFisher), respectively. Both media contained 10% FBS (Gibco), 100 IU ml^−1^ penicillin/streptomycin, 100 µg ml^−1^ amphotericin B (Lonza) and 200 nM L-Glutamine (Gibco). The E0771 and MC38 cancer cells were engineered to express firefly luciferase (Addgene plasmid #19166, gift from Eric Campeau & Paul Kaufman) for bioluminescence imaging.

All cells were grown in a humidified incubator at 37°C, with 5% CO2, and were tested regularly for mycoplasma contamination.

### Animals used

The C57BL/6J-*Gt(ROSA)26Sor^em(CAG-LSL-UPRT-T2A-NLS-sfGFP-IRES-rtTA^*^3^*^-WPRE-pA)1Smoc^* (*R26^LSL-UPRT-HA-sfGFP^*) mice in a C57BL/6 genetic background were generated by inserting CAG-LSL-UPRT^HA^-T2A-NLS-sfGFP-IRES-rtTA3-WPRE-pA expression cassette into ROSA26 locus using a CRISPR– Cas9-mediated genome editing system by Shanghai Model Organisms Center (SMOC).

C57BL/6J (wild type, #000664), *Clec4f ^tdT-Cre^* (#033296), *CD68^EGFP^* (#026827), *Cx3cr1^CreERT^*^2^ (#021160), *Ccr2^GFP^* (#027619), *CD45.1* (#033076) (all from Jackson Laboratory), *H11^CAG-RSR-ZsG-CAG-LSL-tdT^* (denoted as *H11^LSL-tdT^*, #T058308), *H11-CAG-DreERT2* (denoted as *H11^DreERT^*^2^, #T049774), *Ki67^RSR-Cre^* (#T050101), *R26^LSL-DTA^* (#T009408) (all from GemPharmatech), *Tnfrsf11a^Cre^*(#B-EM-009, from Biocytogen), *Id3^flox^* (denoted as *Id3^fl^*, #NM-CKO-2100278) and *R26^LSL-RSR-tdT-DTR^* (#NM-KI-190086, both from SMOC). Genomic DNA was prepared from mouse ear punch with Mouse Genotyping Kit (APExBio) as per manufacturer’s protocol. The wild-type allele was used as the control to distinguish knock-in/knock-out allele. Sequences of all primers used to distinguish knock-in/knock-out allele from wild-type allele were listed in **Supplementary Table 1**.

For retro-orbital injection, the mice were anaesthetized using an isoflurane-based small animal anaesthesia unit. Otherwise, ketamine (100 mg/kg)/xylazine (10 mg/kg) were used for anaesthesia.

6-week-old mice were used unless otherwise specified. No randomization or blinding was used to allocate experimental groups. No statistical methods were used to predetermine sample size.

All animal work was done in accordance with a protocol approved by the Sun Yat-sen University Cancer Center (SYSUCC) Institutional Animal Care and Use Committee.

### Clinical sample collection

6 cases of liver metastasis samples of breast cancer patients (3 cases HER2+ and 3 cases TNBC) were collected from SYSUCC. Paraffin-embedded sections of 3-μm thickness were used for haematoxylin and eosin (HE) staining and immunofluorescence staining. Before immunofluorescence staining, the sections were deparaffinized, and treated with a heat-induced antigen retrieval Tris-EDTA (pH 9.0) solution. After cooling to room temperature, the slides were first washed with running water for 5 min, then washed with 1×PBS twice for 5 min each time.

### Portal vein injection

E0771 cells and MC38 cells were collected from culture with 0.25% trypsin (HyClone). The cells were washed twice with PBS, counted, then resuspended in cold PBS to a concentration of 5×10^6^ cells ml^−1^ and 2×10^6^ cells ml^−1^ for E0771 and MC38, respectively. Mice were anaesthetized, and two 1-inch incisions were made into the shaved abdomen and the peritoneum respectively to reveal the liver. The duodenum was gently pull out to expose the portal vein. The cancer cells were pipetted up and down several times before transferred to 31-gauge needle (BD Bioscience). 50 μl cold PBS were injected into portal vein. The incisions of the peritoneal membrane and skin were closed by absorbable sutures and would clips, respectively. The wound clips were removed on Day 7 post-injection of cancer cells.

### Fat pad injection and subcutaneous injection

E0771 cells and MC38 cells were collected using 0.25% Trypsin-EDTA (Gibco). The cells were washed twice with PBS, counted, and then resuspended in 1:1 solution of PBS and growth factor reduced Matrigel (Corning). For mammary fat pad injection of E0771 cancer cells, mice were anaesthetized, and a small incision was made on the shaved abdomen to reveal mammary gland. 0.5×10^6^ cells in 50 µl ice-cold matrigel/PBS mixture were injected directly into the fat pad. The incision was then closed using wound clips, which were removed on day 7 after injection of cancer cells. For subcutaneous injection of MC38 cancer cells, 0.5×10^6^ cells in 50 µl ice-cold matrigel/PBS mixture were injected into loose shoulder skin on the right site of the mouse.

### Bioluminescent imaging and analysis

Mice were injected retro-orbitally with 1.5 mg of D-luciferin (GoldBio) (15 mg ml^−1^ in PBS). Imaging was completed within 1–2 min after injection, using an IVIS Spectrum Living Imaging system and analysis software v4.2 (Perkin Elmer). For *in vivo* bioluminescence images, photon flux was calculated for each mouse by using a rectangular region of interest encompassing the liver of the mouse in a supine position. For *ex vivo* bioluminescence images, photon flux was calculated for each dissected liver by using a circular region of interest along the edge of a 35-mm cell culture dish.

### Tamoxifen and diphtheria toxin treatment

Tamoxifen (MedChemExpress) was dissolved in corn oil (20 mg ml^−1^) and heated at 37°C for 3 hours. To activate *Cx3cr1^CreERT^*^2^-mediated recombinase, each mouse was injected with Tamoxifen (100 mg/kg body weight) by oral gavage every three days, starting from Day 12 post-intraportal injection, as indicated in **Fig. 2e**. For proliferation tracing experiment, DreERT2-mediated recombinase was introduced by a single dose of tamoxifen gavage (100 mg/kg body weight) on Day 12 post-intraportal injection, as indicated in **Fig. 3f**. For KC specific depletion, DreERT2-mediated recombinase was introduced by 8 doses of tamoxifen gavage (2 mg per dose), as indicated in **Fig. 5d**. A single dose of 200 ng DT (List Laboratories) was intraperitoneally introduced one week after the last Tamoxifen injection. The cancer cells were inoculated into liver through portal vein one day after DT injection.

### Liposomal treatment

For fluorescent labelling or pharmacological depletion of mononuclear phagocytes, 100 μl DiI-labelled liposomes (#F70101F), clodronate liposomes (#F70101A) or PBS-control liposomes ((#F70101CA) were injected intravenously (all at the concentration established by FormuMax), as indicated in **Fig. 3c-h** and **Extended Data Fig. 6a-d**.

### Csf1r inhibitor and dual ICB treatment

The liver metastasis-bearing *Ccr2^GFP/GFP^* mice or littermate control *Ccr2^GFP/WT^* mice were evaluated for metastasis burden using *in vivo* bioluminescence imaging. Total flux between 5×10^5^ to 2×10^6^ radiance (p/sec/cm^2^/sr) were considered as micrometastasis. To inhibit metastasis-associated macrophages, mice were fed with 1200 ppm PLX5622 (TargetMol, formulated in AIN-76A standard chow diet) *ad libitum* for 14 days before ex vivo bioluminescence imaging of liver metastasis burden. To block immune checkpoints Pd1, 100 μg anti-Pd1 antibodies (clone 29F.1A12, BioXcell) were intraperitoneally injected into the metastasis-bearing mice starting from the day of *in vivo* bioluminescence imaging and every 3 days subsequently.

### Liver Dissociation

The mice were euthanized by CO^2^ and subjected to whole-animal perfusion. In brief, an incision was immediately made in the right atrium and 10 ml PBS was slowly injected through the left ventricle. The change in colour of the liver should be observed for the successful perfusion.

After whole body perfusion, the liver metastasis nodules and nearby normal tissues were dissected and briefly cut with scissors. The tissue pieces were transferred to a tissue digestion C-tube (Miltenyi) and incubated in digestion buffer containing 1×RPMI, Collagenase A (1 mg ml^−1^, Sigma, #10103586001), DNase I (0.1 mg ml^−1^, Sigma, #10104159001) and 1% heat-inactivated FBS, and mechanically on a Single-cell Suspension Dissociator (Miltenyi). Briefly, the “Tumor_3” program was run on the Dissociator, followed by a 15-minute incubation at 37°C. Next, the “Tumor_3” program was run again to increase the yield of single cells, followed by a 15-min incubation at 37 °C. After a final run of the “Tumor_3” program, the digestion reaction was filtered through a 70-um cell strainer and was then transferred to 2-ml ice-cold RPMI medium to stop the reaction. Cellular debris and red blood cells were removed by the Debris Removal Solution (Miltenyi) and 1× RBC Lysis Buffer (Biolegend), respectively, as per manufacturer’s instructions.

### Flow Cytometry

Single-cell suspensions were prepared as described in the “**Liver Dissociation**” section. 0.1−0.5×10^6^ cells were incubated for 15 min at 4 °C with anti-mouse Fc-block CD16/32 antibody (clone 93, 1:100) and True-Stain Monocyte Blocker (1:100) (both from Biolegend), in 100 µl FACS staining buffer (1.5% FBS and 0.5% BSA in PBS). Cells were subsequently stained with antibodies in FACS buffer at 4 °C for 25 min. The following antibodies against mouse antigens were used (1 μg ml^−1^): anti-CD45 (clone 30-F11), anti-CD45.1 (clone A20), anti-CD45.2 (clone 104), anti-CD11b (clone M1/70), CD11c (clone N418), anti-F4/80 (clone BM8), anti-F4/80 (clone CI:A3-1), anti-Ly6C (clone HK1.4), anti-Ly6G (clone 1A8), anti-Gr1 (clone RB6-8C5), anti-TCRβ (clone H57-597), anti-CD19 (clone 6D5), anti-NK1.1 (clone PK136), anti-CD4 (clone GK1.5), anti-CD8 (clone 53-6.7), anti-Clec4f (Alexa Fluor 647 conjugated, clone 3E3F9), anti-TIM4 (clone RMT4-54), anti-Pd1 (clone 29F.1A12), anti-Ctla4 (clone UC10-4B9), anti-Tigit (clone 1G9), anti-Lag3 (clone C9B7W), rat IgG2aκ isotype control antibody (clone RTK2758), Armenian hamster IgG isotype control antibody (clone HTK888), mouse IgG1κ isotype control antibody (clone MOPC-21), rat IgG1κ isotype control antibody (clone RTK2071) (all from Biolegend). Cells were analysed or sorted using Beckman CytoFLEX or MoFlo Astios EQ, respectively. All data was analysed using FlowJo v10.

### *In vitro* phagocytosis assay

To trigger apoptosis, E0771 cancer cells were treated with 1 μM Staurosporine (Sigma) for 12 hours. Dead Cell Apoptosis Kit (Thermo Fisher) was used to find optimal time of Staurosporine treatment. The apoptotic cells were stained with CypHer5 (GE Healthcare) and seeded together with normal and metastasis-associated macrophages isolated from *Clec4f ^tdT-Cre^;CD68^EGFP^* mice or *H11^DreERT^*^2^*;Clec4f ^tdT-Cre^;R26^LSL-RSR-tdT-DTR^;Ccr2^GFP/GFP^*mice in a non-tissue culture-treated plate at 1:10 phagocyte to cancer cell ratio for 1 hour, 4 hours and 8 hours. Cells were dissociated from plate with trypsin and the phagocytes were assessed by flow cytometry analysis.

### Immunofluorescence staining

The perfused livers were fixed with 4% paraformaldehyde (PFA) overnight at 4 °C. After being washed with PBS three times, the fixed tissues were incubated in 30% sucrose in PBS (w/v) overnight, and frozen in optimum cutting temperature gel (Tissue-Tek; Sakura). Sections of 6-μm in thickness were cut on Leica CM1850 cryotome.

Slides were blocked in PBS with 10% normal donkey serum (Jackson Laboratory), 1% BSA, and 0.3% Triton X-100 for 1 h at 20 °C. Primary antibodies were incubated overnight at 4 °C in the staining solution (5% normal donkey serum, 0.5% BSA and 0.3% Triton X-100). Secondary antibodies were added into the staining solution and incubated for 1 h at 20 °C. Nuclei were stained with Hoechst 33258 (10 µg·ml−1, Thermo Fisher), for 2 min at 20°C. Excess antibodies were removed by washing for 5 min in PBS with 0.5% Tween-20. Nuclei were counter-stained with 1 μg ml^−1^ DAPI for 3 min at room temperature. To reduce the background, TrueVIEW Autofluorescence Quenching Kit (Vector Labs) was used as per manufacturer’s protocol.

Primary antibodies included: anti-Clec4f (ThermoFisher, #PA5-47396, 1:500), anti-Clec4f (Alexa Fluor 647 conjugated, Biolegend, clone 3E3F9), anti-F4/80 (abcam, #ab6640, 1:200), anti-GFP (GFP-1020, Aves Labs, 1:1000), anti-Glutamine Synthetase (abcam, #73593, 1:1000), anti-HA Tag (Bethyl, #A190-138A, 1:500), anti-HA Tag (abcam, #ab9110, 1:500), anti-CD31 (SinoBiological, #50408-T16, 1:2000), anti-IBA1 (abcam, #ab178847, 1:3000), anti-TIM4 (R&D, #AD2929, 1:100), anti-tdT (abcam, #ab167453, 1:500), anti-tdT (arigo, #ARG55724, 1:500), anti-VE cadherin (R&D, #AF1002, 1:100), anti-Ki67 (CTS, #12202), anti-hHB-EGF (DTR, R&D, #AF259, 1:100), anti-Pd1 (R&D, #AF1021, 1:100), anti-CD8α (R&D, #MAB116, 1:100).

Secondary antibodies included: Alexa-488 donkey anti-chicken, Cy3 donkey anti-rabbit, Alexa-647 donkey anti-rat (1:800, all from Jackson ImmunoResearch). Stained sections were visualized by a Zeiss LSM 980 confocal microscope or scanned with Zeiss Axioscan.Z1. ImageJ was used to manually count the number of macrophages and automatic quantify the number of nuclear foci.

### Proinflammatory chemokines detection

Proinflammatory chemokines of normal liver and liver micrometastasis were detected using LEGENDplex™ Mouse Proinflammatory Chemokine Panel (BioLegend, #740451). Briefly, small liver metastatic lesions (1-2 mm size) and adjacent normal liver tissues were harvested 14 days post portal vein injection, and were kept in PBS containing 25mM EDTA and protease inhibitor (cOmplete tablets, Roche). The tissues were then lysed with a tissue homogenizer, and centrifuged at 12,000 rpm for 30 minutes at 4°C. The supernatant was collected and mixed with Assay buffer in a v-bottom plate. The mixture was subsequently incubated with APC-conjugated capture beads, biotinylated capture antibodies, and PE-conjugated streptavidin according to the manufacturer’s instructions. Fluorescence intensity was recorded by a CytoFLEX flow cytometer (Beckman Coulter). Data was further analysed using LEGENDplex™ Data Analysis Software Suite, version 2023-02-15.

### Liver clearing, immunolabeling and three-dimensional imaging

Tissue clearing was performed with Tissue Clearing Kit (Cat# NH210701-FS, Nuohai Life Science). Briefly, after whole-animal perfusion, livers with metastasis nodules were fixed in ice-cold 4% PFA overnight at 4°C. After 3 times of washes in PBS for 2 hours at RT, livers were immersed in 1/2-diluted Solution 1 (S1) at 37°C for 6 hours, and subsequently in 100% S1 for 3 days at 37°C for rapid de-lipidation and decolorization. Then, after 3 times of washes in PBS for 2 hours at RT, livers were immersed in 1/2-diluted Solution 2 (S2) at 37°C for 6 hours, and subsequently in 100% S2 for 3 days at 37°C for further de-lipidation and decolorization. After de-lipidation, the livers were permeabilized in PBS with 0.3% Triton X-100 at 37°C for 1 hour and were blocked in PBS with 10% normal donkey serum, 1% BSA, and 0.3% Triton X-100 at 37°C for 1 day. Primary antibodies were incubated at 37°C in staining solution (5% normal donkey serum, 0.5% BSA and 0.3% Triton X-100) for 2 days. The livers were washed in PBS with 0.3% Triton X-100 at RT for 2 hours and were incubated with second antibody in staining solution at 37°C for 2 days. After 3 times of washes in PBS for 2 hours at RT, livers were immersed in 1/2-diluted Solution 3 (S3) at 37°C for 6 hours, and subsequently in 100% S3 at 37°C for refractive index (RI) matching, until the tissues become transparent. The cleared tissue was imaged by LiTone XL Light-Sheet Microscope (Light Innovation Technology) for 3D imaging. The magnification is 10×; the numerical aperture of detection lens is 0.6; the spatial resolution is 0.58 μm × 0.58 μm; the field view is 6.2 mm × 3.3 mm. The laser channels are 561 nm and 647 nm for detection of red fluorescent protein tdT and Alexa Fluor 647 conjugated anti-Clec4f antibody, respectively. The original data was processed by LitScan 3.2.0 (Light Innovation Technology).

### *In situ* RNA profiling

The in situ RNA profiling with SLAM-ITseq was conducted as previously described^14^. Briefly, 4-Thiouracil (4tU, Sigma, #440736) was dissolved in DMSO at 200 mg ml^−1^ and was then diluted in corn oil at a 1:4 ratio (50 mg ml^−1^). The solution was vigorously shaken and intraperitoneally injected into *Clec4f ^tdT-Cre^*;*R26^LSL-UPRT-HA-sfGFP^;Ccr2^GFP/GFP^*mice bearing liver metastasis. Liver metastasis tissues and nearby normal tissues were harvested 4 hours after the injection of 4tU without whole animal perfusion. The collected tissues were briefly washed with PBS and immediately transferred to the lysis buffer for RNA extraction (Vazyme, #RC112). RNA was dissolved in H^2^O with 1mM DTT (Sigma, #646563) to prevent oxidation of thiol group in 4tU. 1 μg RNA was reacted with Iodoacetamide (IAA, Sigma, #I1149) buffer (10 mM IAA and 500 mM NaH^2^PO^4^) at 50°C for 15 min to alkylate the thiol group. The RNA from the alkylation reaction was then purified using RNA Clean & Concentrator (Zymo, #R1015), and 0.5 μg RNA was used as input for strand-specific library Prep Kit (Vazyme, NR604). RNA-seq library preparation was conducted following manufacturer’s instructions with the 14 PCR cycles for library amplification. The multiplexed libraries were sequenced using NovaSeq 6000 (Illumina) for pair-end 150 cycles.

A customized version of SLAM-seq data processing pipeline was used for data processing. Briefly, the raw RNA reads were trimmed with Trimmomatic (v0.39) with the parameters ‘SLIDINGWINDOW:5:20 MINLEN:36’. The reverse complemented version of the trimmed R2 reads was further generated by Seqtk (v1.3). Given that Cre^+^ cells incorporated 4tU into nascent RNA and resulted in many T>C conversions after alkylation reaction, the mutation-tolerant aligner NextGenMap (v0.5.5) was used to align the trimmed R1 and reverse complemented R2 reads to mouse reference genome GRCm38 with options ‘-t 8 -n 1 --strata --bam --slam-seq 2’. Samblaster (v0.1.26), Samtools (v1.9), and Slamdunk (v0.4.3; options: filter -mq 2 -mi 0.8 -nm - 1) were used to sort, index, and filter the resulting bam files, respectively. The bam file was further converted to mpileup file by Samtools for downstream analysis with options ‘-B -A -- output-QNAME’. Varscan (v2.3.9) was used to call variants into a vcf file with options ‘mpileup2snp --strand-filter 1 --min-var-feq 0.2 --min-coverage 10 --variants 1’. The mpileup file was screened along with the vcf file to identify reads with at least two T>C base conversion events. Such base converted reads were extracted from the raw read sets by Seqtk to run the mapping process again with NextGenMap for downstream visualization in IGV (v2.16.0) and further analysis. After transcript quantification with RSEM (v1.3.3), R package DESeq2 (v1.34.0) was used to normalise the count matrix and to identify differentially expressed genes.

Enrichment of pathways pertinent to normal or metastasis-associated KCs was identified by R package clusterProfiler (v4.2.2).

### Single cell RNA sequencing

Single cell suspensions (2 ×10^6^ cells ml^−1^ in ice-cold PBS) were loaded onto microwell chip using the Singeron Matrix Single Cell Processing System. Barcoding Beads were subsequently collected from the microwell chip, followed by reverse transcription of the mRNA captured by the Barcoding Beads and to obtain cDNA for PCR amplification. The amplified cDNA was then fragmented and indexed with sequencing adapters. The single cell RNA sequencing (scRNA-seq) libraries were constructed according to the protocol of the GEXSCOPE scRNA Library Kits (Singleron Biotechnologies). Individual libraries were diluted to 4 nM, pooled, and then sequenced on Novaseq 6000 (Illumina) with 150 bp paired end reads.

Singeron scRNA-seq analysis pipeline CeleScope (v1.9.0) was used to pre-process the raw reads and to generate gene expression matrix. Cells with more than 30,000 unique molecular identifier (UMI) countings were filtered out. The filtered count matrices were converted to Seurat (v4.3) objects. Cells expressing less than 200 genes or with more than 20% mitochondrial contents were excluded for downstream analysis. Filtered data were then log normalised and scaled using ‘NormalizeData’ and ‘ScaleData’, respectively. Cell–cell variation due to UMI counts and percent mitochondrial reads regressed out. Top 2000 variable genes were selected by “FindVariableFeautres” for principal component analysis (PCA). The top 20 principal components were used for clustering and dimensional reduction at a resolution of 2.0. The cell type identity of each cluster was determined with the expression of canonical markers based on the annotation of SynEcoSys database (https://singleron.bio/product/detail-25.html). Uniform Manifold Approximation and Projection (UMAP) algorithm was applied to visualize cells in a two-dimensional space. Heatmaps, dot plots, violin plots displaying the expression of markers used to identify each cell type were generated by ‘DoHeatma’, ‘DotPlot’. ‘Vlnplot’, respectively.

Pseudo-time trajectory analysis was performed with Monocle3^ref35^ using default parameters. In brief, the Seurat object was first converted to Monocle CellDataSet. The raw count expression matrix was then pre-processed to normalise data. And top 50 principal components were used to capture variated genes across all the cells in the data set. After dimension reduction with UMAP method, the cells were put in order by how much progress they’ve made by ‘learn_graph’ function. Finally, the cells were ordered in pseudotime along a trajectory and visualized by ‘plot_cell’ function.

For cell indexing and epitopes profiling, single cell suspension was incubated with the cell hashtag antibodies (unique DNA barcodes listed in **Supplementary Table 2**) and the cocktail of 119 unique cell surface antibodies (Biolegend, #199901), as per manufacturer’s protocol.

To trace Kupffer cell-derived LMAMs, an exogenous gene sequence, IRES-WPRE, was appended to the ends of the mouse reference genome FASTA file, accompanied by a GTF file detailing gene annotations. The reference index of this custom genome was constructed using the Celescope ‘mkref’ command for subsequent scRNA-seq analysis.

The Kupffer cell (KC) gene signature and the scar-associated macrophage (SAM) gene signature^24^ were used to illustrate the phenotypic plasticity of different macrophage populations in liver metastasis. The KC genes were enriched in the macrophage cluster of normal mouse liver, while SAM genes were highly expressed in macrophages from chronic carbon tetrachloride (CCl^4^)-induced fibrotic livers.

### Epigenetic profiling

The liver metastasis and nearby normal tissues of *H11^DreERT^*^2^*;Clec4f ^tdT-Cre^;R26^LSL-RSR-tdT-DTR^;Ccr2^GFP/GFP^* mice were dissociated as described in the “**Liver Dissociation**” section. The CD45 Microbeads (RWD Life Science, #K1304) were used to enriched immune cells from single cell suspension through magnetic cell separation. Live cells (HelixNIR^−^) expressing F4/80 and tdT were purified by FACS sorting.

Transposase accessibility and transcription factor binding were profiled using an ATAC-seq kit (#N248) and a CUT-Tag kit (#N259, both from NovoProtein), respectively, as according to manufacturer’s instruction. The following antibodies were used for CUT-Tag assay: anti-Histone H3 acetyl K27 antibody (H3K27ac, abcam, ab177178), anti-Histon H3 tri-methyl K27 (H3K27me3, CTS, #9733), anti-PU.1 (abcam, ab227835), and goat anti-rabbit IgG secondary antibodies (abcam, #ab7085). The PCR cycles was optimized for different factors (ATAC-seq, 10 cycles; PU.1, 16 cycles; H3K27me3, 14 cycles; H3K27Ac: 14 cycles). The multiplexed libraries were sequenced using Novaseq 6000 (Illumina) for pair-end 150 cycles.

Low-quality reads were filtered as described in the “***In situ* RNA profiling**” section and were aligned to the mouse genome GRCm38 with bowtie2 (v2.4.5) with the parameters ‘--local --very-sensitive --no-unal --no-mixed --no-discordant --phred33 -I 10 -X 2000’. Picard (v2.19.0) and bedtools (v2.30.0) were used to remove duplicated reads and blacklist-region reads, respectively. The clean BAM files were converted to bigwig files with deeptools (v3.5.3) bamCoverage using the parameters ‘--effectiveGenomeSize 2652783500 --normalizeUsing RPKM --extendReads 200 --binSize 50 --outFileFormat bigwig’. The bigwig files were uploaded to local tracks of WashU Epigenome Browser (https://epigenomegateway.wustl.edu/) to visualize different histone modification markers and transcription factors simultaneously at specific genome loci.

In addition to genome browser tracks of individual genes, the bigwig files were also used to calculate scores per genome regions with deeptools computeMatrix using parameters ‘scale-regions -S --upstream 3000 --regionBodyLength 5000 --downstream 3000 –skipZeros -- outFileName genes_scaled.mat.gz’. The mitochondria genes in KC signature genes were excluded to eliminate background noises. This intermediate file was used to generate heatmap and profile plot with ‘plotHeatmp’ and ‘plotProfile’, respectively.

Peak calling was performed using macs3 (3.0.0b3) with default parameters. ‘macs3 -q 0.05’ was used for ATAC-seq and CUT&Tag-seq of histone modification antibodies, and ‘macs3 -q 0.25’ was used for CUT&Tag-seq of the transcription factors. The output bed files were used as input for motif analysis, which was conducted by homer (v4.11) findMotifsGenome with default parameters.

### Statistics

In most *in vivo* experiments, group sizes were determined based on the results of preliminary experiments and no statistical method was used to predetermine sample size. Experiments were repeated at least twice independently, and the data are combined and presented. For the dot plots of IF staining results, smaller dots without an outline are values from individual fields (∼0.4 mm^2^ fields), and circles that are outlined represent mean values taken over 4−5 fields from the same mouse, unless otherwise noted. The statistical tests were performed by comparing the individual animals. All *in vitro* experiments were repeated independently for three times (batches). The *P* values in these *in vitro* experiments were determined based on biological replicates.

## Data and code availability

Raw sequencing reads for **Fig. 5g** and **Extended Data Fig. 5a-e** are deposited in the China National Center for Bioinformation (CNCB) under accession number PRJCA024187, and are publicly available as of the date of publication. Processed Seurat objects of scRNA-seq assays for **Fig. 1d** and **Fig. 4b**, and analysis code related to SLAM-ITseq, scRNA-seq, CUT-Tag and ATAC-seq will be uploaded to Github (https://github.com/lintian0616/KC_plasticity). All other data supporting the findings of this study are available from the corresponding author on reasonable request.

## Supporting information

Primers for genotyping

Hastag antibodies for scRNA-seq

3D imaging of cleared liver metastasis

3D imaging of cleared liver metastasis

## Acknowledgment

We thank S-C.Zhang for critically reading the manuscript. L.Tian is supported by the National Natural Science Foundation (NSFC) of China (Grant No. 82173278), the Science and Technology Projects in Guangzhou (Grant No. 202201011364) and Young Talents Program of SYSUCC (Grant No. YTP-SYSUCC-0045); Y-X.Shi and L.Tian are supported by National Key Research and Development Program of China (Grant No. 2021YFE0206300); H-Q.Ju and L.Tian are supported by the Guangdong Basic and Applied Basic Research Foundation (Grant No. 2023B1515040030); H-Y.Huang is supported by Fostering Program for NSFC Young Applicants (Tulip Talent Training Program) of SYSUCC (Grant No. TTP-SYSUCC-202406).

## Author contributions

H-H.Y and L.T conceived the idea. H-H.Y, Y-Z.C, and L.T performed most experiments and wrote the manuscript. X-N.Z and L.T performed epigenetic profiling. L.T performed most bioinformatic analyses. J-X.Y built a custom pipeline to process pair-end fastq data for SLAM-ITseq assay. Y-X.S collected liver metastasis tissue samples from breast cancer patients. H.Q Ju provided reagents and expertise and participated in data analysis. All co-authors have seen and approved the manuscripts.

