## Extended Data Figures 1-6 for "Availability of an inflammatory macrophage niche drives phenotypic and functional alterations in Kupffer cells"

**EXTENDED DATA FIGURE LEGENDS**

**Extended Data Figure 1. Kupffer cells phagocyte apoptotic and liver tumor cells**

**a).** Schematic diagram of the *in vitro* co-culture experiment. The immune cells were enriched using CD45-magnetic beads (MACS, magnetic-activated cell sorting). E0771 cells were treated with Staurosporine (STS) for 12 hours to induce apoptosis, followed by the labeling with CypHer5E pH-sensitive fluorescent dyes.

**b,c).** Flow cytometry gating (**b**) and quantification (**c**) of N-macs (tdT$^{+}$ EGFP$^{+}$) and LMAMs (tdT$^{-}$ EGFP$^{+}$) from *Clec4f* $^{tdT\text{-}Cre}$*;CD68*$^{EGFP}$ mice (n=3).

**d).** Schematic of the experimental design (dpi, days post portal vain injection).

**e).** IF staining for Clec4f, tdT and F4/80. White arrowhead indicating the tumor-derived material tdT engulfed by KC (scale bar, 50 μm).

Mean ± s.e.m. shown. *P* values were calculated by comparing individual animals using two-tailed paired (**c**) Student's *t*-test.

**Extended Data Figure 2. Lineage Tracing of KCs using embryonic macrophage specific Cre recombinase mice**

**a).** Schematic showing *Tnfrsf11a*$^{Cre}$*;R26*$^{LSL\text{-}UPRT\text{-}HA\text{-}sfGFP}$ mice for tracing embryonic-derived macrophages.

**b,c).** IF staining (**b**) and flow cytometry (**c**) quantification of HA tag$^{+}$ or sfGFP$^{+}$ fraction in N-macs and LMAMs (n=6 for IF staining; scale bar, 50 μm; n=5 for flow cytometry).

**d).** Schematic of experimental design. Parabiosis of *CD45.2*$^{+/+}$*;Tnfrsf11a*$^{Cre}$*;R26*$^{LSL\text{-}UPRT\text{-}HA\text{-}sfGFP}$ mouse surgically paired with a *CD45.1*$^{+/+}$*;CD68*$^{EGFP}$ partner for 2 weeks prior to the portal vein inoculation of E0771 tumor cells into *CD45.1*$^{+/+}$*;CD68*$^{EGFP}$ mouse.

**e).** Representative IF staining of HA tag$^{+}$ macrophages in LvMet (scale bar, 50 μm).

**f).** Flow cytometry gating and quantification of bloodborne CD45.2$^{+}$sfGFP$^{-}$ mo-macs (number of parabionts=5).

**Extended Data Figure 3. Quantification of inflammatory chemokines in liver metastasis and nearby normal tissues**

**a).** Schematic of experimental design and representative images of small metastasis lesions for chemokine quantification (scale bar, 2 mm).

**b).** Box-whisker plots showing the enhanced chemokine level in metastatic tissues. The upper and lower hinges correspond to the first and third quartiles, and the upper and lower whiskers are highest and lowest values that are within 1.5×IQR (interquartile range) of the hinge (n=5). *P* values were calculated by comparing individual animals using two-tailed paired Student's *t*-test.

**Extended Data Figure 4. WPRE as a heritable tag for Cre$^{+}$ cells**

**a).** Schematic of experimental design. The two subsets of NK1.1+ cells, NK cells (Cre$^{+}$ sfGFP$^{+}$ TCR$^{-}$) and NKT cells (Cre$^{-}$ sfGFP$^{-}$ TCR$^{+}$), were sorted for single cell TCR sequencing (scTCR-seq).

**b).** Integrated genome viewer plot showing transcripts mapping to the WPRE.

**c).** Summary of hashtag antibodies and number of cells yielded after pre-processing steps in the scTCR-seq experiment.

**d).** UMAP embedding of Cre+ NK cells and Cre$^{-}$ NKT cells from MC38 LvMet and nearby normal tissues.

**e,f).** Feature plot (**e**) and bar plot (**f**) showing the distribution and quantification Cre$^{+}$ cells based on WPRE expression.

**Extended Data Figure 5. *In Situ* transcriptome profiling of KCs using SLAM-ITseq**

**a).** Schematic of experimental design showing the incorporation of 4-Thiouracil (4tU) into nascent RNA.

**b).** Bioinformatic analysis pipeline (left) and representative genome view plot showing the T->C and A->G point mutations in 4tU-incorporated nascent RNA.

**c).** Principal component analysis (PCA) of KCs based on SLAM-ITseq of normal liver and LvMet of MC38 and E0771 (n=3 per group).

**d,e).** Dot plot (**d**) and heatmap (**e**) showing the enriched Gene Ontology (GO) terms in KC-derived LMAMs (KIF, kinesin family members; SMC, structural maintenance of chromosomes genes; CDC/CDK, cell division cycle genes/cyclin dependent kinase genes; ANAPC, anaphase promoting complex subunit genes).

**f).** Representative IF staining and quantification of Ki67$^{+}$ fraction in HA tag$^{+}$ KC-derived LMAMs (n=6; scale bar, 50 μm).

Mean ± s.e.m. shown. *P* values were calculated by comparing individual animals using two-tailed paired (**f**) Mann–Whitney *U*-test.

**Extended Data Figure 6. Pharmacological depletion and genetic impairment of KCs**

**a).** Schematic of experimental design (CL, clodronate liposomes; PVi, portal vein injection; BLI, bioluminescence imaging; HE, haematoxylin-eosin staining).

**b-d).** Quantification of metastatic outgrowth based on normalised bioluminescence values (**b**), the numbers of surface metastasis nodules (**c**) and HE staining (**d**) in CL-treated mice (n=6 for MC38; n=7 for E0771) and PBS control liposome treated mice (n=6 for MC38; n=7 for E0771).
**e,f).** Quantification of metastatic outgrowth based on normalised bioluminescence values (**e**) and the numbers of surface metastasis nodules (**f**) in $Tnfrsf11a^{Cre};Id3^{fl/fl}$ mice (n=9 for MC38; n=7 for E0771) and $Id3^{fl/fl}$ littermates (n=8 for MC38; n=6 for E0771).
The *ex vivo* bioluminescence value was normalised to the *in vivo* bioluminescence value obtained immediately after portal vein injection (day 0). Mean ± s.e.m. shown. *P* values were calculated by comparing individual animals using two-tailed unpaired (**b**-**d**, **e**, **f**) Mann–Whitney *U*-test.

**Extended Data Figure 7. Dual blockade of monocyte infiltration and macrophage proliferation in primary tumors**

**a).** Representative IF staining of IBA1$^+$ tumor-associated myeloid cells in orthotopic E0771 breast tumor (scale bar, 50 μm).
**b,c).** Flow cytometry quantification of tumor-infiltrating monocytes (**b**) and tumor-associated macrophages (TAMs) (**c**) in $Ccr2^{GFP/GFP}$ mice fed with PLX5622 diet (n=7 for MC38, n=5 for E0771) or control chow (n=5 for MC38, n=7 for E0771; subQ, subcutaneous), and in $Ccr2^{GFP/WT}$ littermates fed with Csf1r inhibitor PLX5622 diet (n=4 for MC38, n=6 for E0771) or control chow (n=5 for MC38, n=5 for E0771).
Mean ± s.e.m. shown. *P* values were calculated by comparing individual animals using two-tailed unpaired (**b**, **c**) Mann–Whitney *U*-test.

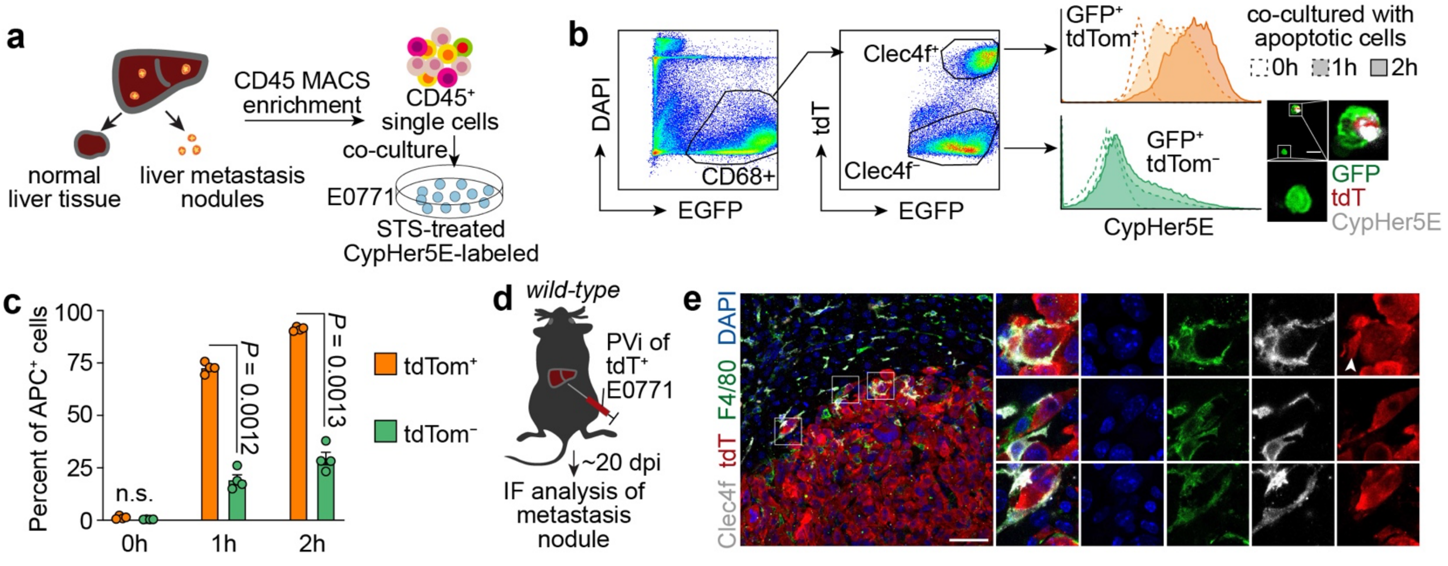


**Extended Data Fig 1. Kupffer cells phagocyte apoptotic and liver tumor cells**

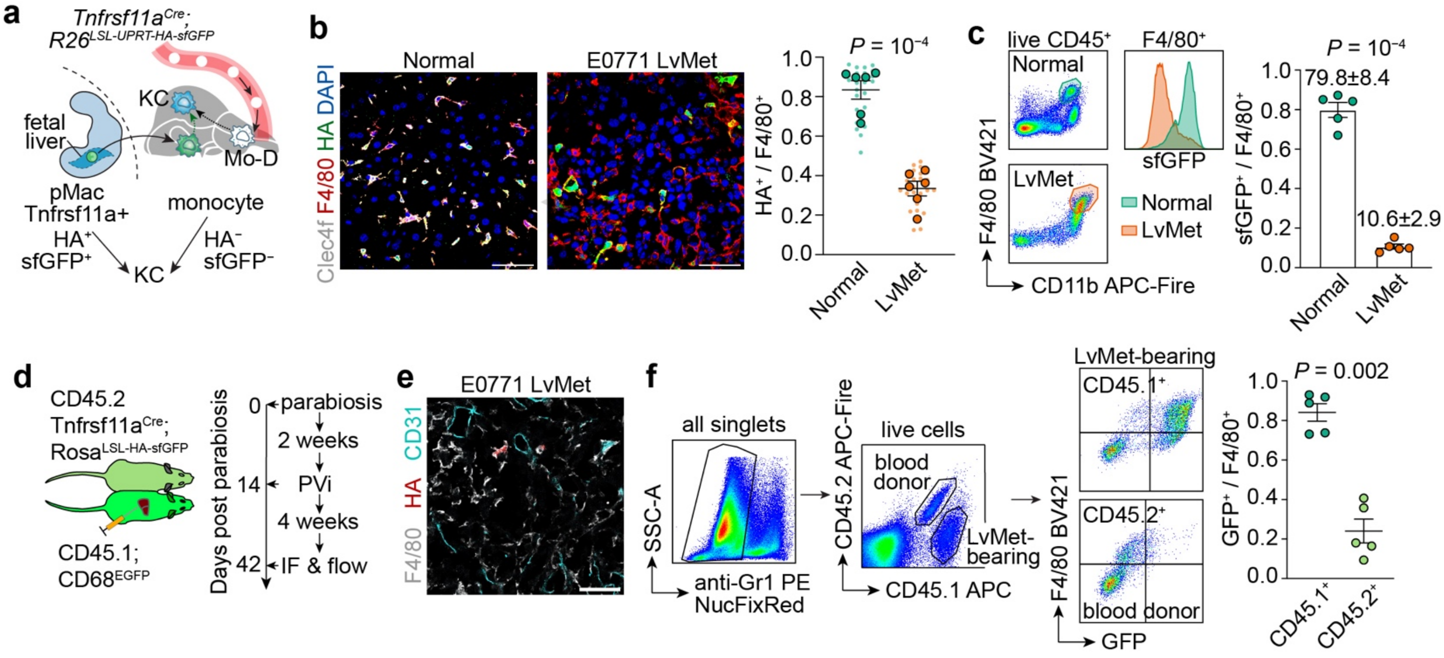


**Extended Data Figure 2. Lineage Tracing of KCs using embryonic macrophage specific Cre recombinase mice**

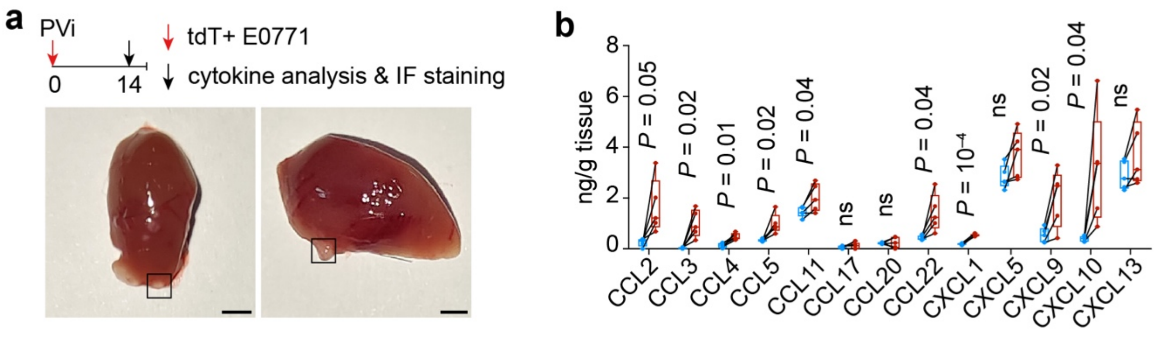


**Extended Data Figure 3. Quantification of inflammatory chemokines in liver metastasis and nearby normal tissues**

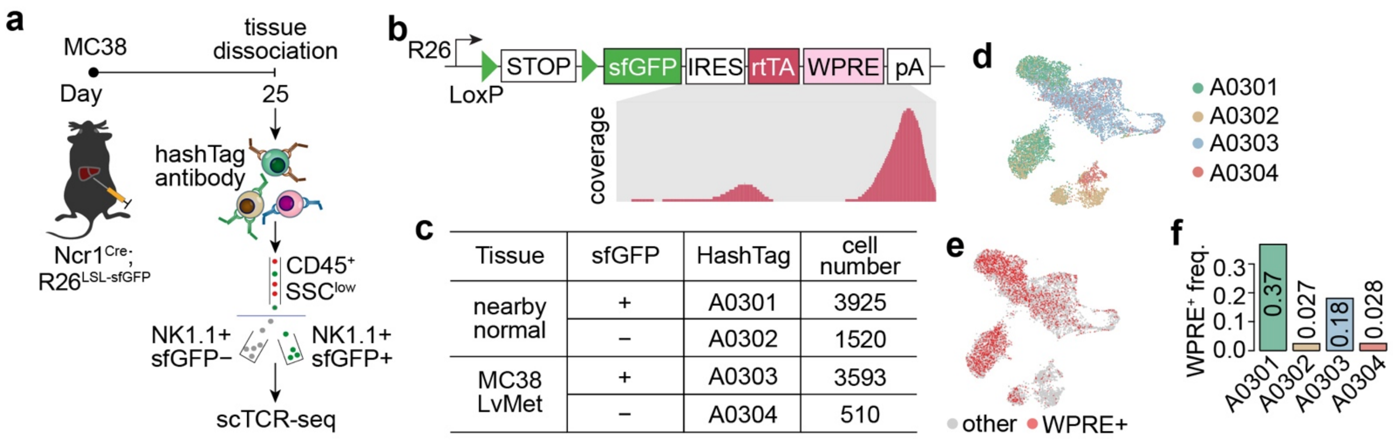


### Extended Data Figure 4. WPRE as a heritable tag for Cre⁺ cells

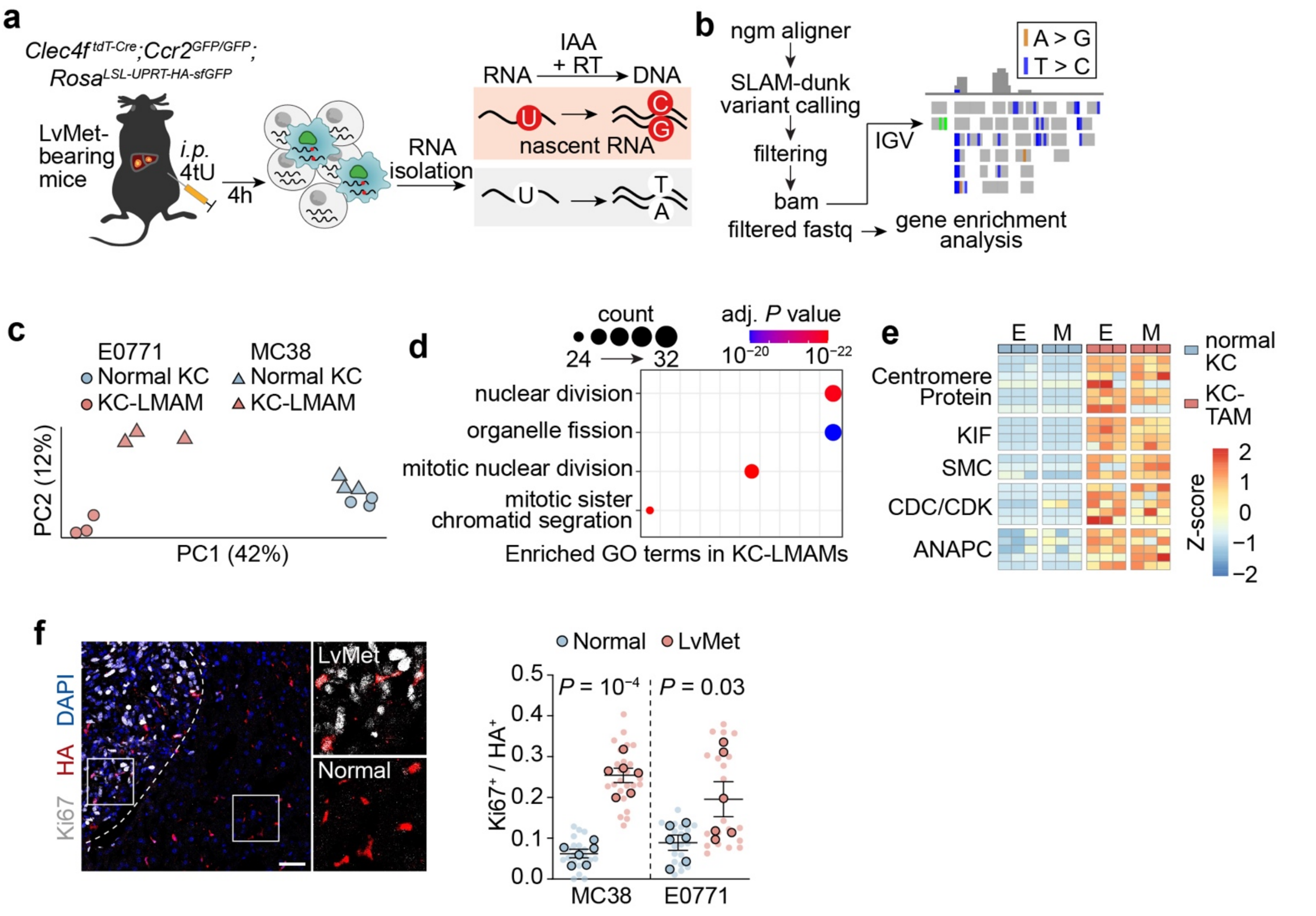


**Extended Data Figure 5. *In Situ* transcriptome profiling of KCs using SLAM-ITseq**

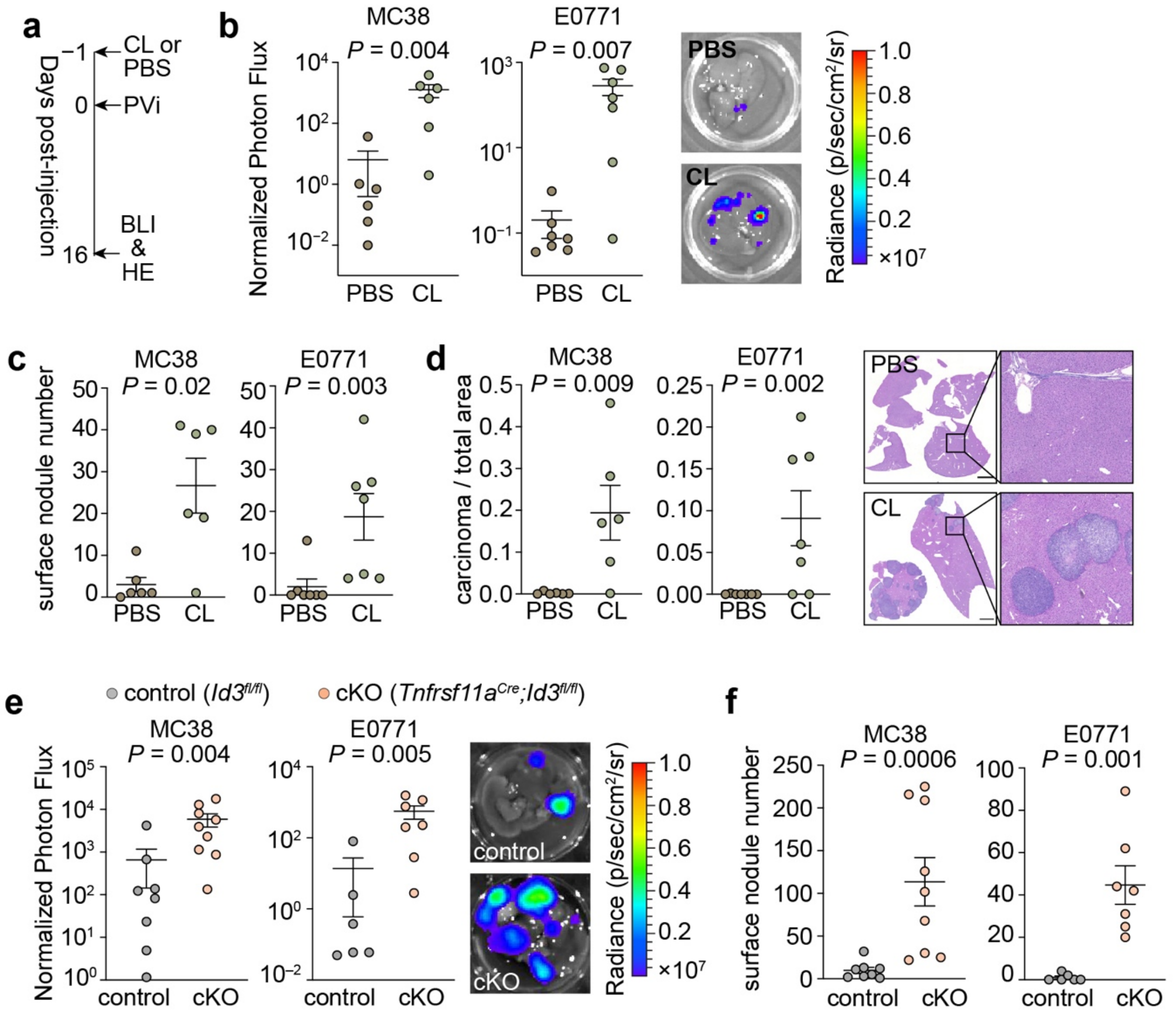


**Extended Data Figure 6. Pharmacological depletion and genetic impairment of KCs**

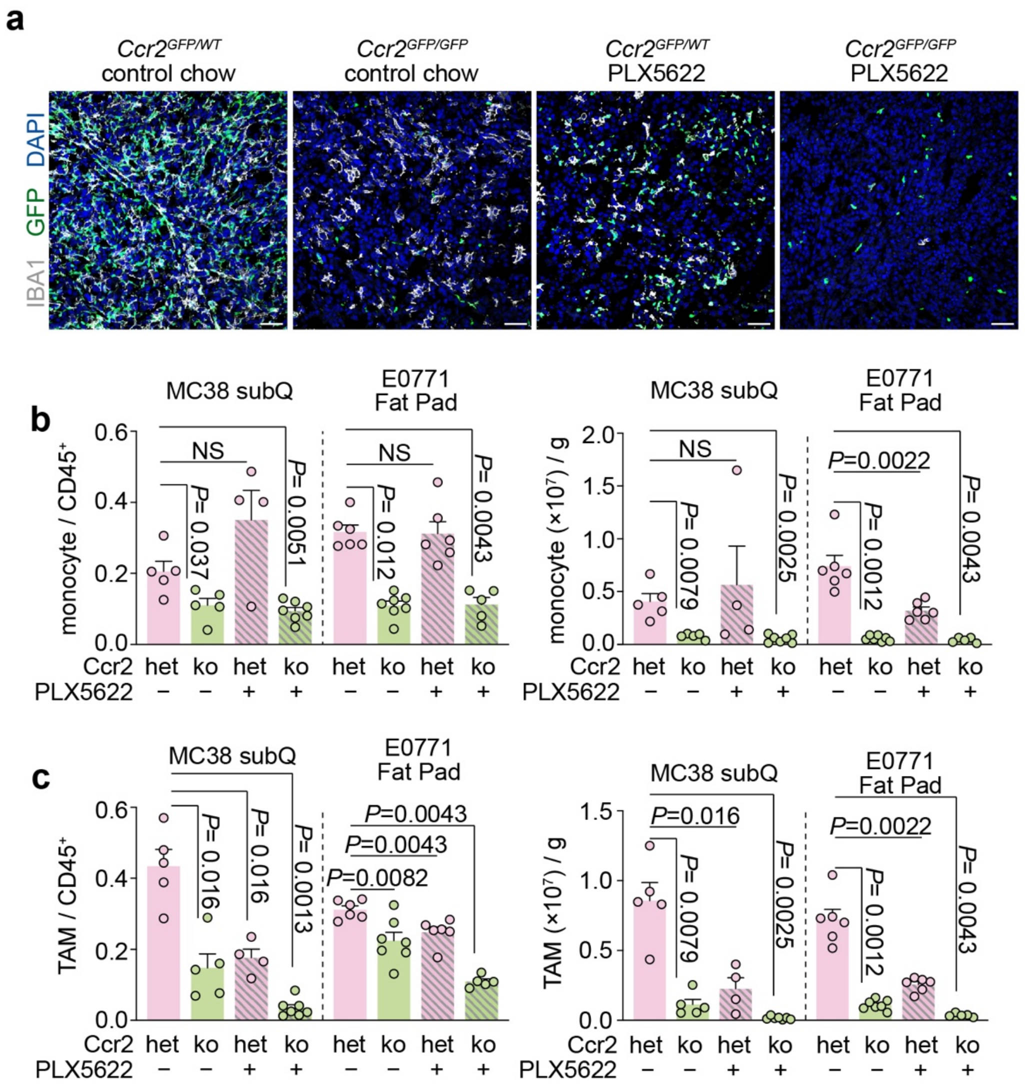


**Extended Data Figure 7. Dual blockade of monocyte infiltration and macrophage proliferation in primary tumors**
