## Supplementary material for "Availability of an inflammatory macrophage niche drives phenotypic and functional alterations in Kupffer cells": Primers for genotyping

### Refer to Fig. 1a

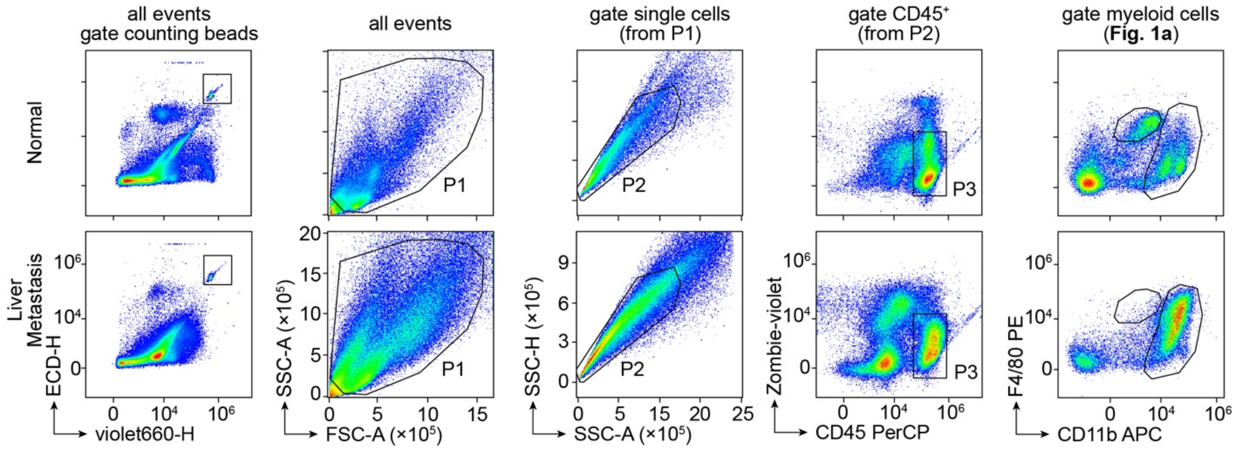


### Refer to Fig. 1b,c (gating single cells as shown in Fig. 1a)

**Myeloid cells (DC-P7, Neutrophils-P4, Monocytes-P5, CD11b$^{high}$ Mac-P8, CD11b$^{int}$ Mac-P2)**

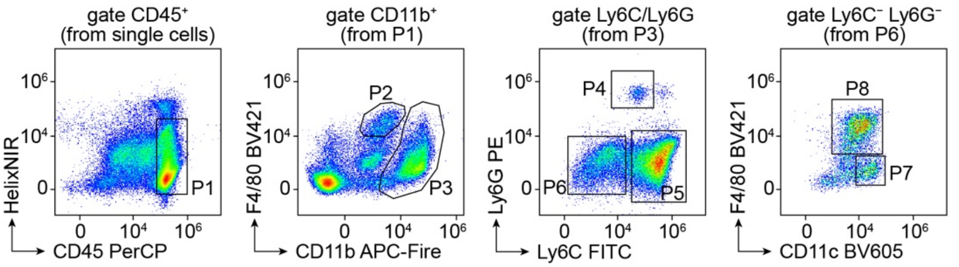


**Lymphoid cells (NKT-P5, NK-P4, CD8⁺ T-P7, CD4⁺ T-P8, B cells-P2), gating CD45⁺ cells as shown in "Myeloid cells"**

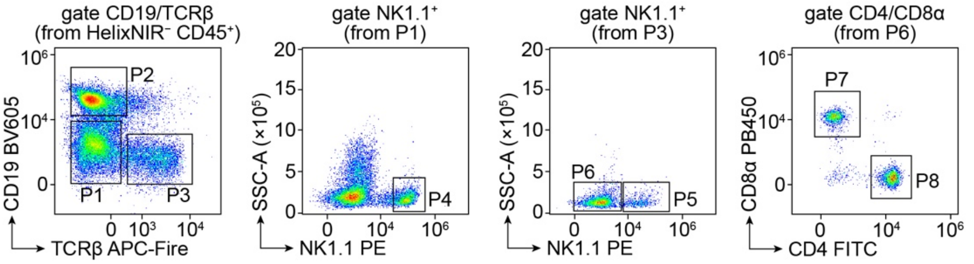


### Refer to Fig. 1g,h (gating single cells as shown in Fig. 1a)

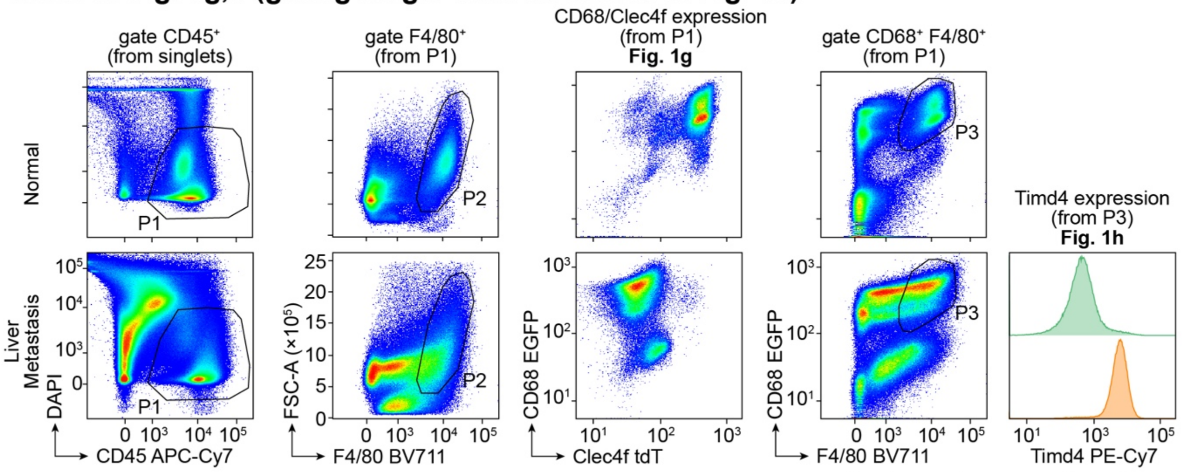


**Gating strategy for Fig. 1**

### Refer to Fig. 2c

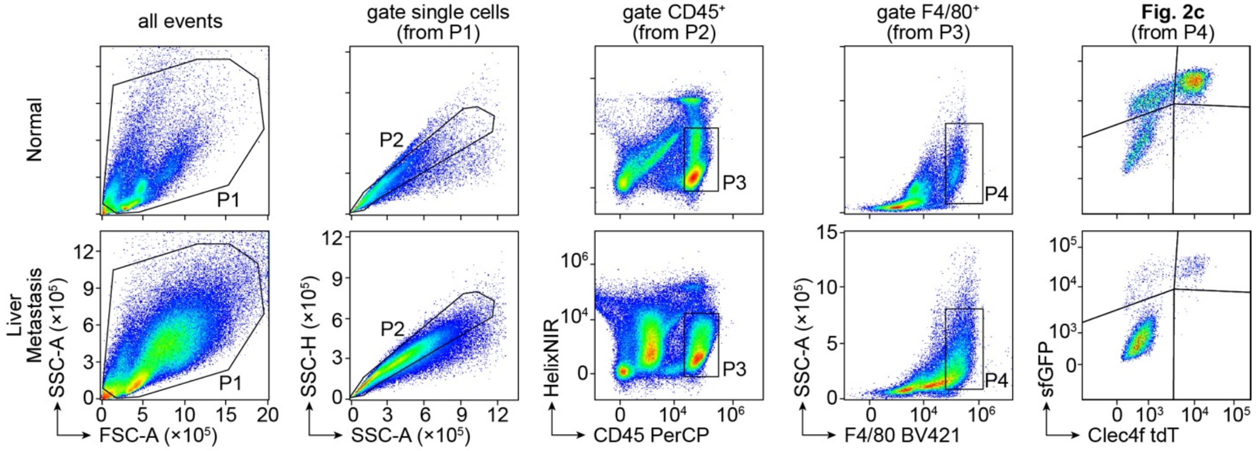


### Refer to Fig. 2f (gating F4/80$^+$ cells as shown in Fig. 2c)

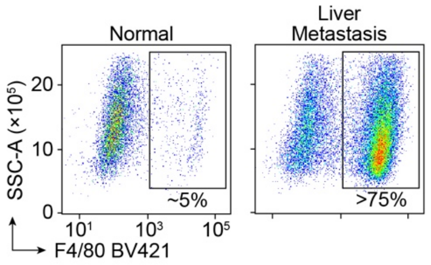


### Refer to Fig. 2j (gating CD45$^+$ cells as shown in Fig. 2c)

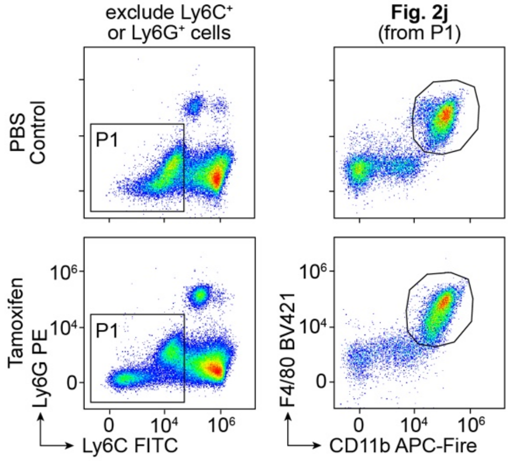


### Refer to Fig. 2l (gating CD45$^+$ cells as shown in Fig. 2c)

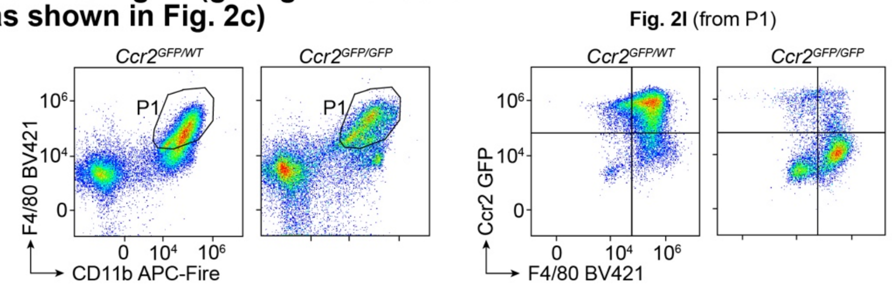


**Gating strategy for Fig. 2**

### Refer to Fig. 3c,f

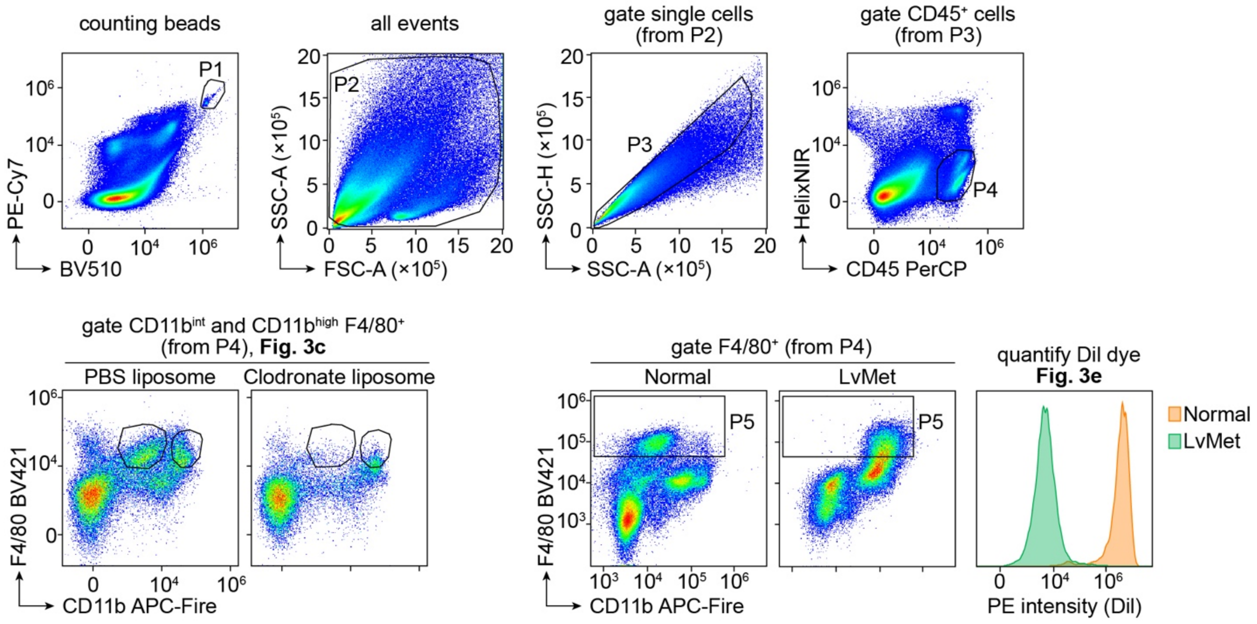


### Refer to Fig. 3h (gating CD45⁺ cells as shown in Fig. 3c,f)

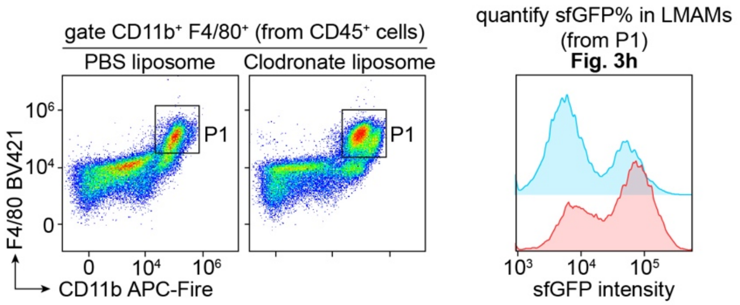


**Gating strategy for Fig. 3**

### Refer to Fig. 5c

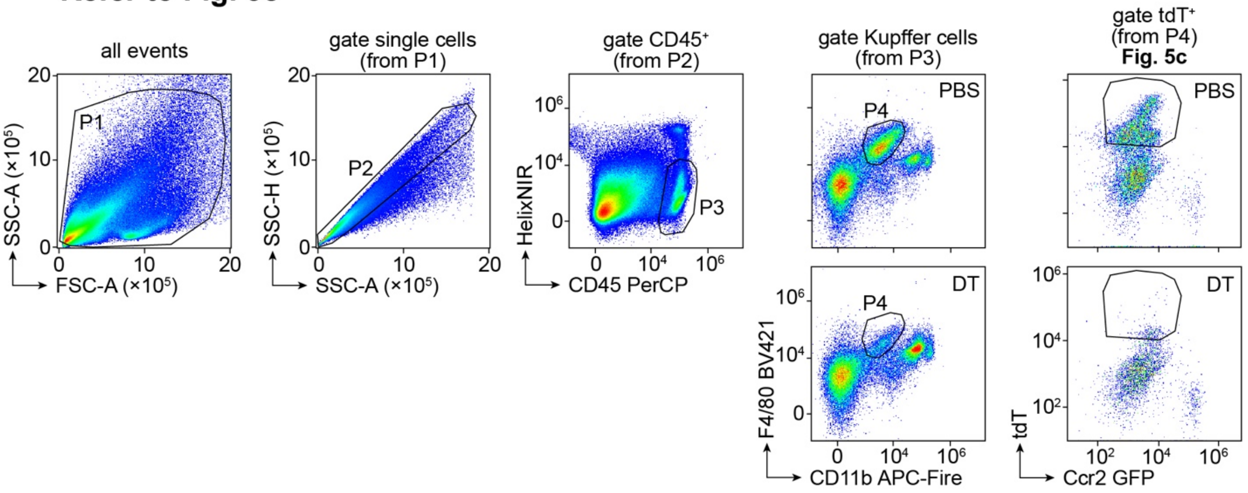


### Refer to Fig. 5f (gating single cells as shown in Fig. 5c)

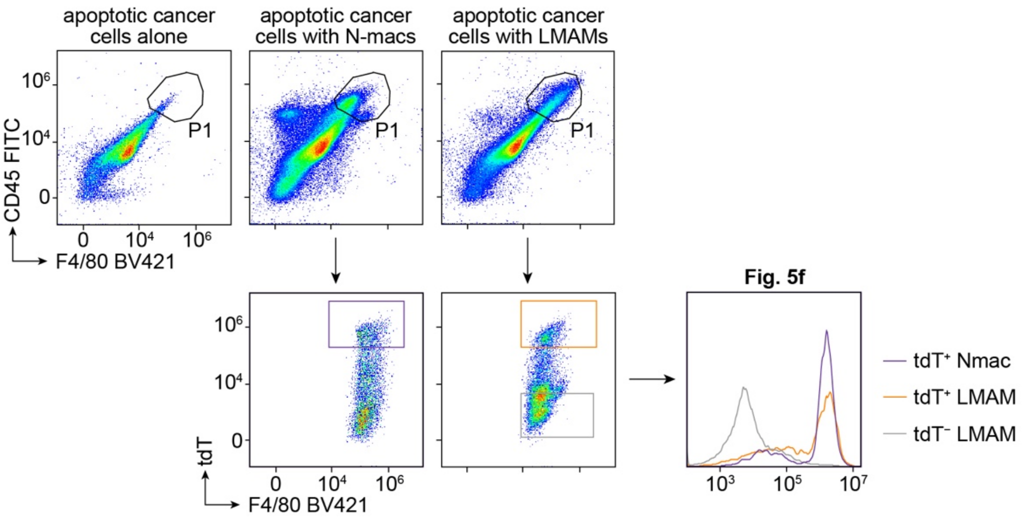


**Gating strategy for Fig. 5**

### Refer to Fig. 6a,b and Extended Data Fig. 7b,c

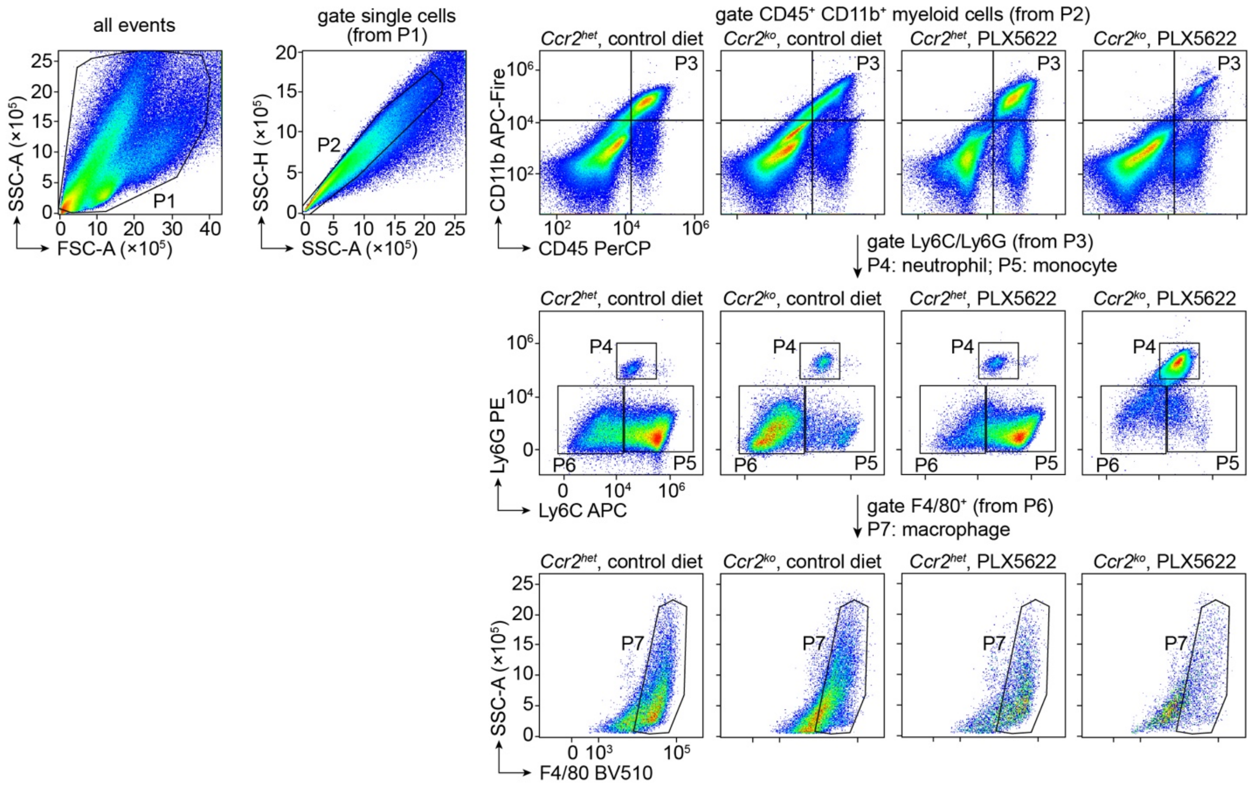


### Refer to Fig. 6e (gating single cells as shown in Fig. 6a,b)

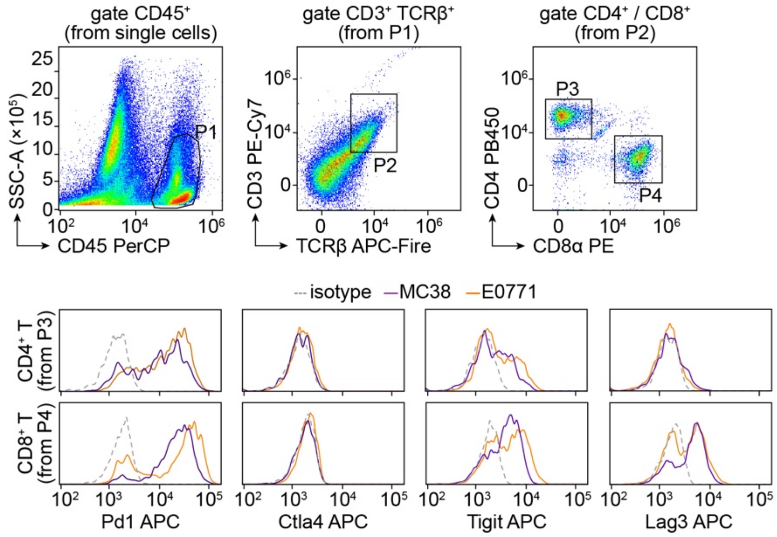


**Gating strategy for Fig. 6**

### Refer to Extended Data Fig. 1b,c

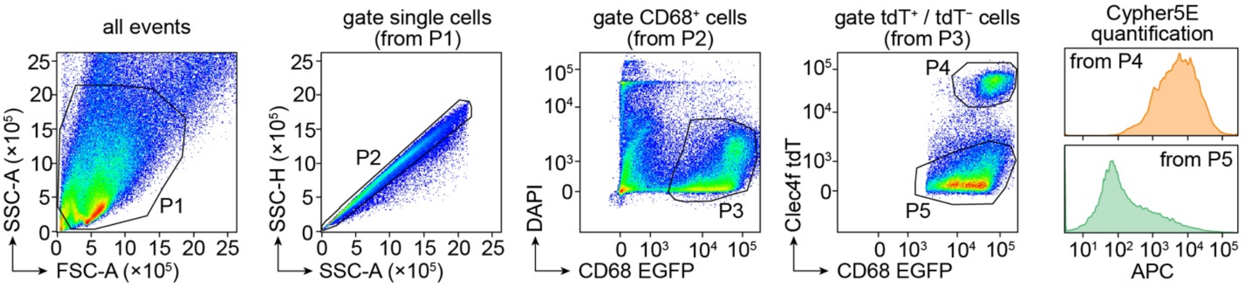


**Gating strategy for Extended Data Fig. 1**

### Refer to Extended Data Fig. 2c

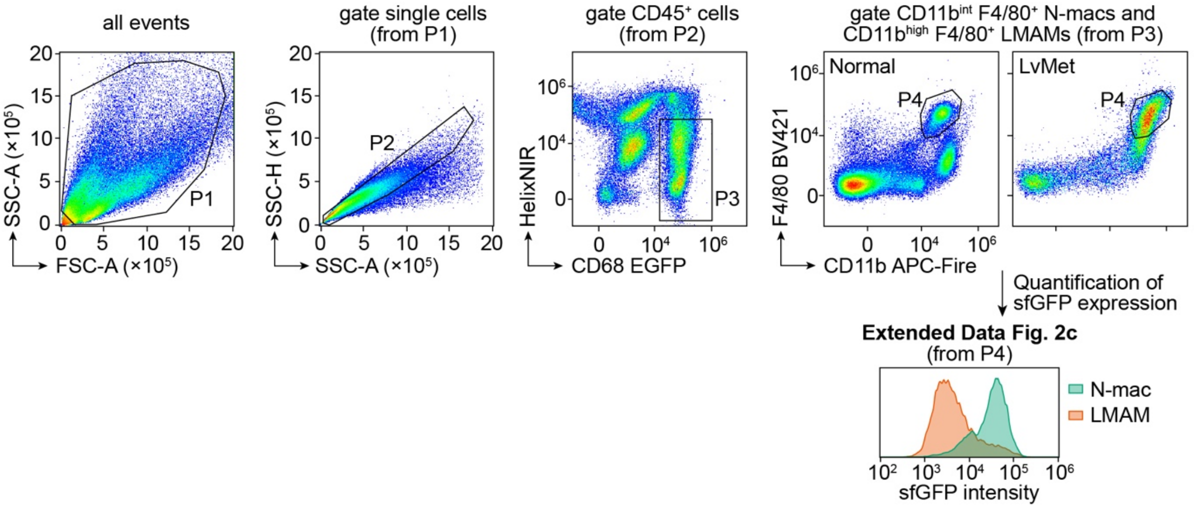


### Refer to Extended Data Fig. 2f (gating single cells as shown in Extended Data Fig. 2c)

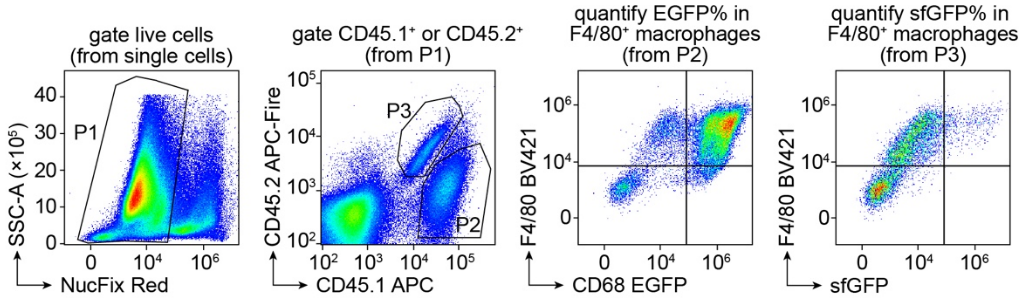


**Gating strategy for Extended Data Fig. 2**
