## Supplementary material for "Availability of an inflammatory macrophage niche drives phenotypic and functional alterations in Kupffer cells": Hastag antibodies for scRNA-seq

**Supplementary Table 1. Sequence of Primers for Genotyping**

| MouseStrain | Primers (5' -> 3') |
|---|---|
| Clec4f-tdT-Cre | common_F: CAAGAAGTCCACAGGGTGGT |
| | WT_R: GAAAGACCCAAGGGAAGGAG |
| | KI_R: ACACCGGCCTTATTCCAAG |
| CD68-EGFP | tg_F: CGGCTCTGTGAATGACAATG |
| | tg_R: TTGGCAGTTGTGGCAAGTAG |
| Cx3cr1-CreERT2 | common_F: AAGACTCACGTGGACCTGCT |
| | WT_R: AGGATGTTGACTTCCGAGTTG |
| | KI_R: CGGTTATTCAACTTGCACCA |
| Ccr2$^{GFP}$ | common_F: AATAATCATTTTGTTCTCTGACCAC |
| | WT_R: ACAGCATGAACAATAGCCAAGT |
| | KI_R: CTGAACTTGTGGCCGTTTAC |
| CD45.1 | PCR_F: GCAAGACTGTACTCCATGAGT |
| | PCR_R: ATTTATGGCGGATCACTGAGG |
| | seq: CAGAGTTGAGAGGGTTCACAT |
| LSL-tdTom | WT_F: AGTCTTTCCCTTGCCTCTGCT |
| | WT_R: GGGTCTTCCACCTTTCTTCAG |
| | KI_F: GGGCAGTCTGGTACTTCCAAGCT |
| | KI_R: TCAATGGAAAGTCCCTATTGGCGT |
| CAG-DreERT2 | WT_F: CAGCAAAACCTGGCTGTGGATC |
| | KI_F: CCTCCTCTCCTGACTACTCCCAGTC |
| | common_R: ATGAGCCACCATGTGGGTGTC |
| Ki67-RSR-Cre | common_F: CAGAGCTAACTTGCGCTGACTGGA |
| | WT_R: CCCGATTCCATTTGGAAGCTC |
| | KI_R: TAGAAGGCACAGTCGATCCTCTAG |
| LSL-DTA | common_F: CCCAAAGTCGCTCTGAGTTGTTA |
| | WT_R: TCGGGTGAGCATGTCTTTAATCT |
| | KI_R: TAGTCTCGTGCAGATGGACAGCA |
| Tnfrsf11a-Cre | common_F: CCTGTGCAGGAGGAGACGCT |
| | WT_R: GAGGTGCACAGTGGCTAGCT |
| | KI_R: TGGTGCACAGTCAGCAGGTTG |
| Id3$^{flox}$ | common_F: CCGTGGTATCTGGGTTTGCT |
| | WT_R: CTGGGGACAAGATTAACACAGG |
| | KI_R: GTCTGGGGACAAGATTAAATAACTTCG |
| LSL-RSR-tdT-DTR | WT_F: TCAGATTCTTTTATAGGGGACACA |
| | WT_R: TAAAGGCCACTCAATGCTCACTAA |
| | KI_F: ATGAAGCTGCTGCCGTCGG |
| | KI_R: TCAGTGGGAATTAGTCATGCCCAA |
| LSL-UPRT$^{HA}$-sfGFP | common_F: TCAGATTCTTTTATAGGGGACACA |
| | WT_R: TAAAGGCCACTCAATGCTCACTAA |
| | KI_R: AAGTAGCAAAAGAGGAAAGAGAGACA |

**Supplementary Table 2. Hashtag Antibodies Used for scRNA-seq (Fig. 3)**

| MouseModel | Tissue | Sorted Cell Number | Hashtag Antibody (5'->3') | Dilution |
|---|---|---|---|---|
| Clec4f$^{Cre\text{-}tdT}$; CCR2$^{GFP/WT}$; LSL-UPRT$^{HA}$-sfGFP-WPRE$^{ki/wt}$ | Normal Liver | 19,979 | TotalSeq™-A0301 ACCCACCAGTAAGAC | 1:100 |
| | MC38 Liver Metastasis | 29,838 | TotalSeq™-A0302 GGTCGAGAGCATTCA | |
| Clec4f$^{Cre\text{-}tdT}$; CCR2$^{GFP/GFP}$; LSL-UPRT$^{HA}$-sfGFP-WPRE$^{ki/wt}$ | Normal Liver | 20,018 | TotalSeq™-A0303 CTTGCCGCATGTCAT | |
| | MC38 Liver Metastasis | 34,452 | TotalSeq™-A0304 AAAGCATTCTTCACG | |
